# Differential roles of putative arginine fingers of AAA^+^ ATPases Rvb1 and Rvb2

**DOI:** 10.1101/2024.05.13.593962

**Authors:** Jennifer L. Warnock, Jacob A. Ball, Saman M. Najmi, Mina Henes, Amanda Vazquez, Sohail Koshnevis, Hans-Joachim Wieden, Graeme L. Conn, Homa Ghalei

**Author notes:** Current address: WuXi Biologics, New Jersey, USA. Current address: Teleflex Incorporated, Pennsylvania, USA.

## Abstract

The evolutionarily conserved AAA^+^ ATPases Rvb1 and Rvb2 proteins form a heteromeric complex (Rvb1/2) required for assembly or remodeling of macromolecular complexes in essential cellular processes ranging from chromatin remodeling to ribosome biogenesis. Rvb1 and Rvb2 have a high degree of sequence and structural similarity, and both contain the classical features of ATPases of their clade, including an N-terminal AAA^+^ subdomain with the Walker A motif, an insertion domain that typically interacts with various binding partners, and a C-terminal AAA^+^ subdomain containing a Walker B motif, the Sensor I and II motifs, and an arginine finger. In this study, we find that despite the high degree of structural similarity, Rvb1 and Rvb2 have distinct active sites that impact their activities and regulation within the Rvb1/2 complex. Using a combination of biochemical and genetic approaches, we show that replacing the homologous arginine fingers of Rvb1 and Rvb2 with different amino acids not only has distinct effects on the catalytic activity of the complex, but also impacts cell growth, and the Rvb1/2 interactions with binding partners. Using molecular dynamics simulations, we find that changes near the active site of Rvb1 and Rvb2 cause long-range effects on the protein dynamics in the insertion domain, suggesting a molecular basis for how enzymatic activity within the catalytic site of ATP hydrolysis can be relayed to other domains of the Rvb1/2 complex to modulate its function. Further, we show the impact that the arginine finger variants have on snoRNP biogenesis and validate the findings from molecular dynamics simulations using a targeted genetic screen. Together, our results reveal new aspects of the regulation of the Rvb1/2 complex by identifying a relay of long-range molecular communication from the ATPase active site of the complex to the binding site of cofactors. Most importantly, our findings suggest that despite high similarity and cooperation within the same protein complex, the two proteins have evolved with unique properties critical for the regulation and function of the Rvb1/2 complex.

**Significance:** AAA ATPases constitute a large family of proteins involved in various essential cellular functions in living organisms in all kingdoms of life. Members of this family typically form homo or hetero multimers that convert the energy from ATP hydrolysis to mechanical work. How the conserved features of AAA ATPases relay the energy from ATP hydrolysis to other functional domains of the complex remains largely unknown. Here, using arginine finger variants of Rvb1 and Rvb2, two evolutionarily conserved closely related AAA^+^ ATPases that form a heterohexameric complex, we reveal how individual protomers in a heteromeric complex can uniquely contribute to the overall function of the complex and how changes in the ATP binding site can be relayed to distal functional domains.

## Introduction

The <u>A</u>TPases <u>a</u>ssociated with diverse cellular <u>a</u>ctivities, termed AAA^+^ ATPases (1), comprise a large class of evolutionarily conserved enzymes involved in diverse essential cellular processes, ranging from DNA replication and damage repair to protein remodeling and degradation to ribosome biogenesis (2–4). AAA^+^ ATPases are identified based on their structural similarities and classified into different clades based on their unique structural features (2, 5). AAA^+^ ATPases are often found as multimers that function as molecular motors powering mechanical work by hydrolyzing ATP to exert conformational changes in other proteins. They can also act as molecular switches or as scaffolds where they work as part of a larger macromolecular machine (3). AAA^+^ ATPases contain several key functional motifs that aid in binding and hydrolysis of ATP. The Walker A, Walker B, Sensor I, Sensor II motifs and arginine fingers are highly conserved motifs that are essential for the ATPase activity (6). The conserved arginine finger is often inserted into the active site of the AAA^+^ ATPases from a neighboring protomer in the multimeric complex to accelerate hydrolysis of the bound ATP (3, 4, 6, 7). The positive charge on the guanidium group of the arginine finger is proposed to stabilize the negative charge evolving on the transition state of ATP-hydrolysis during catalysis (6, 8, 9). Thus, mutations resulting in the loss of the arginine fingers often result in the abolishment of catalytic activity (6).

Yeast Rvb1 and Rvb2 (human RUVBL1 and RUVBL2, respectively) are essential AAA^+^ ATPases that are involved in the assembly of several important ribonucleoprotein complexes, including RNA polymerase II, telomerase, and small nuclear and nucleolar ribonucleoprotein complexes (snRNPs and snoRNPs) (10, 11). To carry out their essential functions, Rvb1 and Rvb2 form a multimeric complex. When purified individually, Rvb1 and Rvb2 can form homohexamers that when mixed results in heterohexameric rings (10, 12–15), referred hereafter as Rvb1/2. The Rvb1/2 complex can further assemble into a dodecameric complex, but the physiological role of these larger assemblies remains unclear (7, 12, 15–19). In the molecular structures characterized to date, only the heterohexameric complex is found to interact with the various binding partners of Rvb1/2 (19–27).

Rvb1 and Rvb2 have a high degree of sequence (65%) and structural (RMSD <1.4 Å) similarity. The overall structure of Rvb1 and Rvb2 has three distinct parts: an N-terminal αβα subdomain of the AAA^+^ domain that contains the Walker A motif (Domain I), an insertion domain that typically interacts with various binding partners of Rvb1/2 (Domain II), and a C-terminal AAA^+^ domain that contains the Walker B motif, the Sensor I and II motifs, and the arginine finger (Domain III) (3, 4, 19, 24, 28) (**Figure 1**). Similar to other AAA^+^ ATPases (4, 6), the conformational dynamics of Rvb1/2 changes as the complex binds, hydrolyzes, and releases nucleotides. Based on structural analyses, binding of ATP by Rvb1/2 protomers induces rotation of the AAA^+^ rings relative to one another, and hydrolysis of ATP allows for further rotation of the rings. These rotations are proposed to alter the interactions of the Rvb1/2 complex with DNA, RNA, or protein-binding partners (7). However, the exact consequences of the different steps during the ATP-hydrolysis-dependent cycle remain poorly understood. Specifically, how these events are regulated and coupled with the role of the complex in various cellular processes has yet to be elucidated. The ATPase activity of the Rvb1/2 complex is essential for its biological functions, as many studies have shown that mutations that abolish the ATPase activity lead to cellular growth defects and impact the cellular pathways that Rvb1/2 is involved in (29–36).

**Figure 1.**
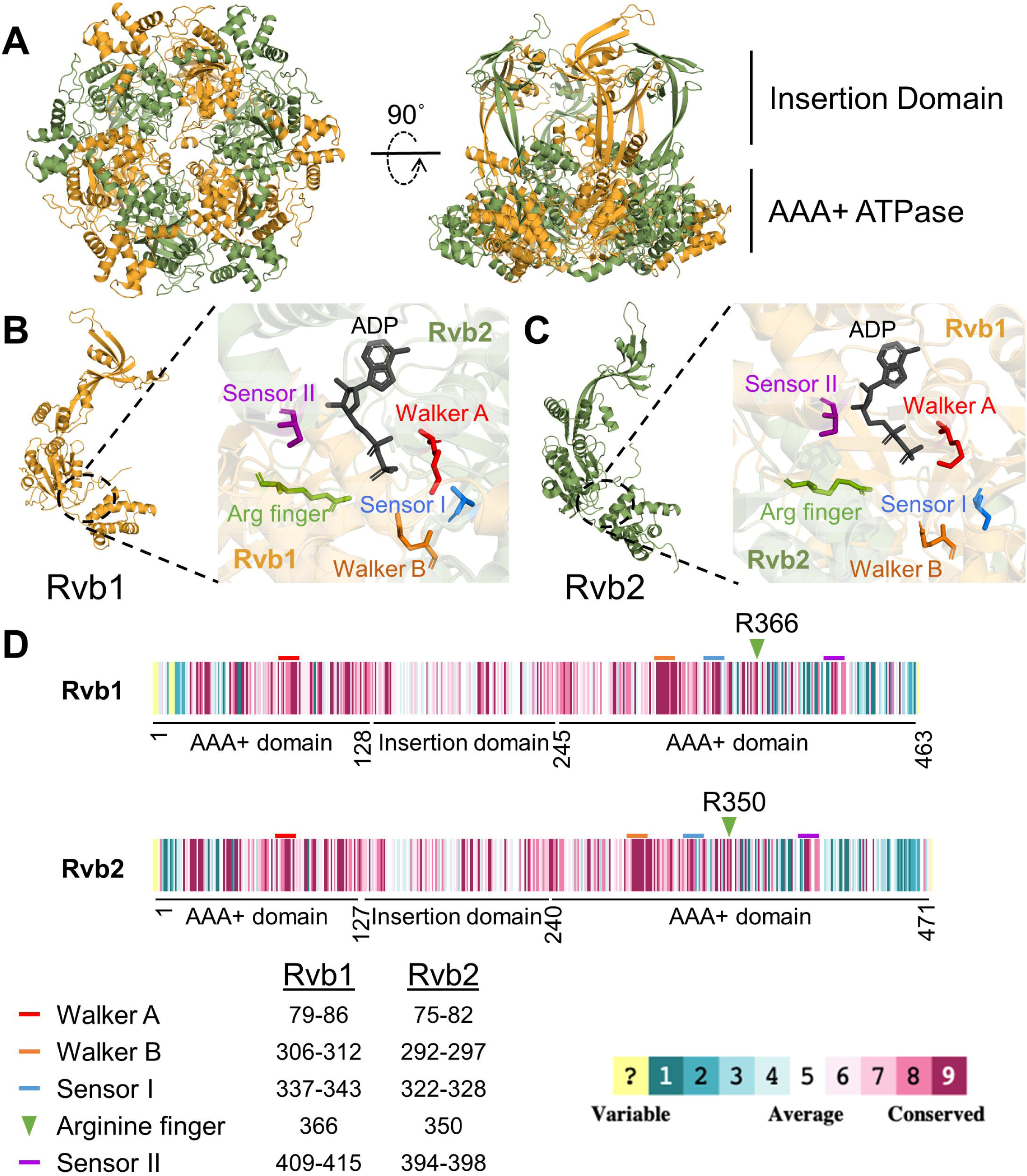
AAA+ ATPases Rvb1 and Rvb2 share a high degree of sequence and structural similarity. **A**) Structure of Rvb1/2 heterohexamer (PDB 6GEJ), highlighting the insertion domain and AAA^+^ ATPase domain. Rvb1 is in orange and Rvb2 is in green. **B-C)** Structure of individual Rvb1 (**B**) and Rvb2 (**C**) protomers and their ATP binding domains (PDB 6GEJ). Key ATPase residues are highlighted, including the conserved trans-arginine fingers (green). Individual subunits colored as in **A**. ADP molecule shown in grey. **D)** Sequence conservation map of yeast Rvb1 and Rvb2 showing the conservation status of the sequences between *S. cerevisiae* and other eukaryotes. Conservation scale indicating level of sequence conservation shown below. Figure made using Consurf. Key ATPase sequence motifs indicated by colored lines or arrows.

Rvb1/2 has been shown to have lower ATPase activity compared to other AAA^+^ ATPases (10, 18, 28, 37, 38). The slow catalytic rate of Rvb1/2 suggests that protein-binding partners or cofactors may be needed *in vivo* to accelerate the catalytic activity. Indeed, to carry out their function, Rvb1/2 heterohexamers typically form complexes with other proteins, such as Tah1 and Pih1, in the context of the R2TP complex (23, 28), the chromatin remodeling INO80 complex (20, 22, 37, 39), the SWR-C complex (24, 27), and the snoRNP biogenesis factors (40, 41). Structural studies have shown that Rvb1/2 associates with these various factors via the DII insertion domain. In the context of R2TP complex, Tah1/Pih1 can stimulate Rvb1/2’s ATPase activity by roughly 1.5-fold when added to reaction mixtures (28). The binding of Rvb1/2 to Ino80 results in 16-fold increased ATPase activity of the complex (37). The rate-limiting step in the reaction of the bacterial homolog of Rvb1/2, RuvB, involves dissociation of ADP, where the arginine finger induces conformational rearrangements necessary for ADP release (42). However, whether this is a general feature of all oligomeric AAA^+^ ATPases remains unknown.

The ATPase activity of Rvb1/2 is essential for proper snoRNP assembly (11, 29–32). Small nucleolar RNAs (snoRNAs) are abundant non-coding RNAs responsible for guiding modifications or processing of other types of RNAs, such as ribosomal RNAs (rRNAs) (43–45). Based on their conserved sequence motifs, snoRNAs are classified into two major groups: box C/D and box H/ACA snoRNAs (46). Rvb1/2 is required for the assembly of snoRNAs from both classes into functional complexes and for maintaining the steady-state levels of the snoRNAs (31, 40, 44, 46). Box C/D snoRNAs assemble with four essential proteins-Snu13, Nop56, Nop58, and the methyltransferase Nop1 (fibrillarin) to form a functional snoRNP complex (11, 46). In addition to Rvb1/2, at least five other assembly factors have been identified to be required for C/D snoRNP assembly in yeast. These include Bcd1, Tah1, Pih1, Rsa1, and Hit1 that interact directly or indirectly with Rvb1/2 during snoRNP assembly (11, 47). How the Rvb1/2’s ATPase activity aids in snoRNP assembly steps remains unknown.

Heere, using *Saccharomyces cerevisiae* as a model organism we address the contribution of putative arginine fingers of Rvb1 and Rvb2 in both the enzymatic mechanism of ATP hydrolysis and the Rvb1/2 complex function. Yeast growth assays reveal that identical changes in arginine fingers of Rvb1 vs Rvb2 have very different biological outcomes. *In vitro* ATPase assays show that alanine substitutions of the arginine fingers of Rvb1 or Rvb2 cause drastically different effects on ATPase activity of the complex, suggesting that albeit being part of the same complex, Rvb1 and Rvb2 differentially impact the overall ATP hydrolysis mechanism. Using molecular dynamics (MD) simulations, we find that changes near the active site of Rvb1/2 propagate long-range changes in the structural dynamics of the insertion domain, suggesting how activity within the catalytic site of ATP hydrolysis can be relayed to other domains to influence the protein’s function. To validate MD simulations data, we took snoRNP biogenesis pathway as an example and assessed the impact from arginine finger variants on the interaction with the Nop58 protein that binds to the insertion domain. Western blot analyses reveal that both arginine finger variants significantly reduce the steady-state levels of Nop58 protein, and overexpression of Nop58 can partially rescue the growth defect of cells expressing the arginine finger variant. Finally, we further validate the differential effect of Rvb1 and Rvb2 arginine variants in response to cellular stressors inducing stress granule formation, where Rvb1/2 is shown to be involved. Together, our results reveal differential contributions of Rvb1 and Rvb2 to the complex and suggest that despite high similarity and cooperation within the same complex, the two proteins have evolved with unique properties that impacts the interaction of their complex with clients.

## Methods

### Serial dilution growth assays

All yeast strains used in this study are listed in **Table S1** and all plasmids in **Table S2**. Yeast strains were grown at 30 °C overnight to saturation and then diluted in sterile water to a concentration of 1 x 10^7^ cells/mL. Ten-fold serial dilutions were then made in sterile water and dilutions were plated on growth media before incubation at the indicated temperatures.

### Expression and purification of Rvb1/2 complex

A pET28a plasmid encoding an MGSS-8xHis-TEV-Rvb2-Rvb1 (kind gift from the Hopfner lab) was transformed into Ros II (DE3) *E. coli* cells. A 2 L culture was grown in 2xYT media and expression was induced with 0.3 mM IPTG at OD_600_ ∼0.5. Cultures were then grown at 18°C for 16-18 hours and harvested by centrifugation at 4,000 x g for 20 minutes at 4°C. Pellets were stored at –80°C until purification. Frozen cells were lysed by thawing the pellet in 30 mL of lysis buffer (30 mM Tris HCl pH 7.5, 300 mM NaCl, 20 mM imidazole, 10% glycerol, 2 mM β-mercaptoethanol (BME)) with added 0.1 mg/mL lysozyme, 1 mM PMSF, 10 µM E64, and 10 µM pepstatin. Cells were incubated in lysis buffer for 15 minutes on ice, and then lysed by sonication for 2 mins total with 3 seconds on, 10 seconds off intervals on the 55% power setting on ice. Lysates were clarified by centrifugation at 18,000 rpm for 45 minutes at 4 °C. The supernatant was applied to 4 mL of packed Ni-NTA resin and incubated while rocking for 45 minutes at 4 °C. The flowthrough was collected and added to an additional 2 mL packed fresh Ni-NTA resin and incubated while rocking for 30 minutes at 4 °C. Ni-NTA resin was then washed twice with 25 mL of lysis buffer, once with an ATP-Mg^2+^ buffer (50 mM KCl, 10 mM MgCl_2_, 4 mM ATP) to release bound chaperones, and twice more with 25 mL of lysis buffer. The protein was eluted with lysis buffer supplemented with 300 mM imidazole. Eluates were pooled, and TEV protease (1 mg/mL) was added to a 1:50 mL ratio. The mixture was dialyzed overnight in a buffer containing 30 mM Tris HCl pH 7.5, 200 mM NaCl, 5% glycerol and 2 mM BME. After overnight dialysis, the protein sample was added to 2 mL packed Ni-NTA resin and reverse Ni-NTA affinity was used to collect cleaved Rvb1/2. This eluate was diluted 1:1 with a no salt buffer (30 mM Tris HCl pH 7.5, 5% glycerol, 2 mM BME) and loaded onto a 5 mL Hi-Trap Q ion exchange column equilibrated with a low-salt buffer (30 mM Tris HCl pH 7.5, 100 mM NaCl, 5% glycerol, 2 mM BME). The protein was eluted by a stepwise gradient starting from 5% low-salt buffer to 30% high-salt buffer (30 mM Tris HCl pH 7.5, 1 M NaCl, 5% glycerol, 2 mM BME) over 10 column volumes (CV), followed by 30% high salt buffer over 5 CVs, and finally 30% to 100% gradient over 10 CVs. The pure fractions were pooled, concentrated and loaded onto a Superose 6 10/300 increase size exclusion chromatography column in a buffer containing 20 mM Tris HCl pH 7.5, 150 mM NaCl, 1 mM DTT. Fractions of high purity that migrated together in a distinct peak away from the column’s void were pooled, concentrated, frozen in liquid nitrogen, and stored at –80 °C.

### Single turnover ATPase assays

The ATPase activity of Rvb1/2 and variants were analyzed under single turnover conditions using radiolabeled ATP [γ-P^32^]. For this assay, 2.5 µM Rvb1/2 and 25 nM radiolabeled ATP were added to the reaction buffer (25 mM Tris HCl pH 7.5, 150 mM KCl, 10 mM MgCl_2_) and incubated at 30 °C for allotted time points. The reactions were quenched by adding KH_2_PO_4_ to a final concentration of 75 mM and spotted onto a thin layer chromatography plate (TLC) and allowed to separate on the plate in a buffer containing 1 M LiCl and 300 mM NaH_2_PO_4_. The TLC plate was then exposed to a phosphor screen and imaged. Bands were quantified using Image Lab software 6.1 (Biorad), and background was subtracted using the rolling disk method.

### Multiple turnover coupled ATPase assay

The ATPase activity of Rvb1/2 and variants were analyzed under multiple turnover conditions using an NADH-coupled assay. For this assay, 100 nM Rvb1/2 was added to a buffer containing 25 mM Tris HCl pH 7.5, 4.5 mM MgCl_2_, 150 mM KCl, 2 mM phosphoenolpyruvate, 10 U/mL lactate dehydrogenase, 10 U/mL pyruvate kinase, 0.18 mM NADH, and ATP at the indicated concentrations. ATPase activity was monitored over 60 minutes at 30°C via the loss in absorbance of NADH measured at 340 nm on a Biotek synergy neo2 microplate reader. The raw data was analyzed by taking the absolute value of the slope and using the Beer-Lambert law to convert the slope to rate in concentration over time. These rates were then compared using one-way ANOVA in Graphpad Prism software.

### Stopped flow ATP association and dissociation assays

Rate constants for Rvb1/2 and respective alanine variants interacting with 2’/3’-O-(N-Methyl-anthraniloyl)-adenosine-5’-triphosphate (mant-ATP) were determined using a KinTek SF-2004 stopped-flow apparatus. The mant group was directly excited at 355 nm and fluorescence emission was monitored after passing through LG-400-F (400 nm long-pass) cutoff filters (Newport Corp.). The resulting fluorescence time courses were fit with a one-exponential function (equation 1) for Rvb1/2 and Rvb1/Rvb2-R350A, or a two-exponential function (equation 2) for Rvb1-R366A/Rvb2 and Rvb1-R366A/Rvb2-R350A, where *k_app_* is the apparent rate constant, *A* is the signal amplitude, *Fl* is the fluorescence at time *t*, and *Fl_∞_*is the final fluorescence signal. The exponential function was selected depending on which equation gave the best fit. Fluorescent data were normalized and averaged (4-6 traces typically). For association experiments, the obtained concentration dependence of *k_app_* was fit with a linear function; the slope represents the bimolecular association rate constant. All experiments were performed in the following buffer: 50 mM Tris-HCl pH 7.5, 150 mM KCl, 10 mM MgCl_2_.

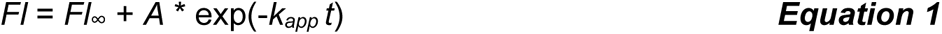

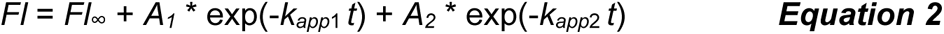

The association rate of mant-ATP to Rvb1/2 was determined by rapidly mixing 25 µL containing 2 µM of the respective protein complex with increasing concentrations (1-6 µM final concentration) of mant-ATP at 20°C. The resulting fluorescent time courses for the Rvb1/2 and Rvb1/Rvb2-R350A were best fit with a one-exponential (equation 1), and a two-exponential function for Rvb1-R366A/Rvb2 and Rvb1-R366A/Rvb2-R350A (equation 2), using TableCurve (Systat Software Inc., USA). To determine the rates of nucleotide dissociation from the respective Rvb1/2 complexes, 2 µM of protein were incubated with 0.5 µM of mant-ADP. Dissociation experiments were then performed by rapidly mixing Rvb1/2*mant-ADP with 100X excess (50 µM final) of unlabeled nucleotide (ATP), at 20°C. Fluorescence time courses were best fit with equation 1. From the obtained rate constants, the equilibrium dissociation constant (*K*_D_) was calculated based on *K*_D_ = *k*_-1_/*k*_1_. The initial binding of mant-ATP to Rvb1/2, resulting in a Rvb1/2•nucleotide complex, is a bimolecular reaction (association rate constant *k*_1_ with units of µM^-1^ s^-1^).

### Western blot analysis

Total cell extracts were prepared by lysis with bead beating in a buffer containing 50 mM Tris-HCl pH 7.5, 150 mM NaCl, 5% glycerol, 2% Triton X-100, 1 mM PMSF, and an EDTA-free protease inhibitor tablet. The lysate was cleared by centrifugation at 10,000 x g for 5 minutes at 4 °C. A BCA assay was performed to measure the total protein concentration of the soluble fraction. Equal total protein concentrations were analyzed on a 12% SDS polyacrylamide gel. Gels were then transferred to nitrocellulose membranes which were incubated in 3% milk with antibodies against Rvb1 and Rvb2 (1:5000, kind gifts from Dr. Walid Houry), GAPDH (1:10,000, Proteintech, cat# HRP-60004), HA (1:1000, Roche, Cat# 11867423001), and FLAG (1:5000, Sigma, cat# F1804). Blots were imaged on a Bio-Rad ChemiDoc MP using horseradish peroxidase-conjugated secondary antibodies and ECL reagent. For quantification, band densities were measured using Image Lab software and normalized to GAPDH loading control or Coomassie stain, where indicated.

### Northern blot analysis

Total cell RNA was isolated from cells grown to A_600_ of ∼0.6, in biological triplicates, using the hot phenol method. RNAs were separated on a 10% Urea polyacrylamide gel and transferred to a Hi-bond nylon membrane. Membranes were probed using oligos listed in **Table S3**, and bands were quantified in Image Lab.

### Measurement of inflection temperature of purified proteins

Purified Rvb1/2 and variants were diluted to 1 µM in a buffer containing 20 mM Tris HCl pH 7.5, 150 mM NaCl and 1 mM DTT. Samples were loaded into glass capillaries and absorbance at 350 nm and 330 nm was measured over increasing temperatures by a Tycho nanotemper instrument.

### Dynamic light scattering analysis of purified proteins

Purified Rvb1/2 and variants were diluted to a concentration of 1.5 µM in a buffer containing 20 mM Tris HCl pH 7.5, 150 mM NaCl, 1 mM DTT. Samples were then loaded into a 384-well plate and analyzed via Dynapro Plate Reader III.

### System Preparation and MD Simulations

A cryo-EM structure of yeast Rvb1/Rvb2 (PDB: 6GEJ) at 3.6 Å resolution was used to generate homology models of Rvb1-R366A/Rvb2, Rvb1/Rvb2-R350A, and Rvb1-R366A/Rvb2-R350A. The resulting homology models were used as starting coordinates for 100 ns MD simulations. All starting structures were prepared using the Protein Preparation Wizard from Schrödinger and MD simulations were carried out in Desmond with the OPLS4 force field. Simulated systems were prepared using the System Building in Desmond. The system was placed within a TIP3P water box. Chloride or sodium ions were used to neutralize the system, and additional sodium and chloride atoms were added to reach a physiological salt concentration of 0.15 M. Before MD production, each solvated system was subjected to series of restrained minimization stages to relax the protein complex. The protein was relaxed in four stages. These stages consisted of successive minimizations with restraints on (i) heavy protein atoms for 200 ps, (ii and iii) protein backbone atoms for 500 ps and 100 ps, respectively, and finally (iv) no restraints for 100 ps. The restraining force constants were 500, 200, and 5 kcal mol^−1^ Å^−2^ for stages (i)−(iii), respectively, and the minimization was done using steepest descent followed by the limited-memory BFGS method to a tolerance of 0.5 kcal mol^−1^ Å^−2^. This tolerance was further reduced to 0.05 kcal mol^−1^ Å^−2^ during unrestrained minimization. Unrestrained MD simulations were carried out in the isothermal−isobaric ensemble using a Langevin thermostat and barostat. The equations of motion were integrated using multiple time steps for the short-range (2 fs) and long-range (6 fs) interactions with a 10 Å cutoff applied for nonbonded interactions. MD simulations for each system were carried out in triplicate at 310 K, with each of the three 100 ns simulations starting with different randomized accelerations. The root-mean-square deviation (RMSD) from the starting structures was assessed and indicates that the simulations reached equilibrium after 10 ns. Only configurations from the equilibrated part of the trajectories were used for subsequent analyses.

### Molecular Dynamics Simulations Analysis

Trajectory analysis was done using Schrödinger Python API and custom scripts. All scripts are publicly available through GitHub. Root-Mean-Square Deviation (RMSD) and Root-Mean-Square Fluctuation (RMSF). RMSD and RMSF were calculated using ***Equations*** 3 and 4, respectively. RMSD plots were used to ensure all simulations reached equilibrium.

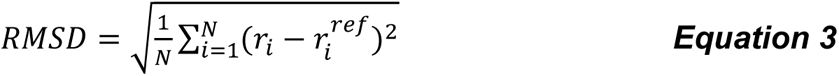

Where N is the number of atoms, 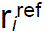 is the coordinates of the i-th atom in the reference structure, r*_i_* is the coordinates of the same atom on the comparison structure.

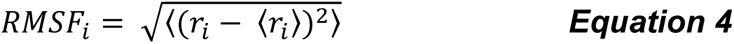

Where r*_i_*is the position of atom *i* at a time point and is the ensemble average.

## Results

### Although highly conserved, the arginine fingers in Rvb1 and Rvb2 differentially impact growth

Rvb1 and Rvb2 are evolutionarily conserved from yeast to man (**Figure S1**). Structural studies of the Rvb1/2 complex have revealed that Rvb1 and Rvb2 form stable heterohexameric rings composed of alternating Rvb1 and Rvb2 protomers (**Figure 1A**) (10, 19, 28). As members of the AAA+ ATPase family, both Rvb1 and Rvb2 subunits contain the conserved motifs common to this family of ATPases, including the Walker A, Walker B, Sensor I, Sensor II motifs and putative arginine fingers (6, 7, 10) (**Figure 1B-C**). Similar to arginine fingers in other ATPases, the arginine fingers of Rvb1/2 are highly conserved across different species (**Figure 1D**) and penetrate from one protomer to contribute to the active site of the neighboring protomer. These arginine fingers are speculated to be involved in the stabilization of the intermediate nucleotide states during ATP hydrolysis (6, 8, 9, 42). On average, the distance of the terminal amino group of arginine fingers from the beta phosphate of the bound nucleotide in the available structures of Rvb1/2 (16, 17, 20–23, 25–27, 39, 48–55) is about 11Å, which is further away than expected if they were involved in assisting the catalysis of the gamma phosphate of the bound nucleotide. The wide range (6-16 Å) suggests the flexibility and interdomain movements of the complex if arginine finger played a role in catalysis or nucleotide binding. Although none of the available structures include the γ-phosphate of the nucleotide, it is evident that without significant structural movement, the arginine fingers appear too far to contribute to nucleotide catalysis in the Rvb1/2 complex. We, therefore, assayed whether variation of the arginine fingers of Rvb1/2 would have any impact on cell viability, as other mutations to the Rvb1/2 ATPase active site have been shown to in the past (31).

To characterize the contribution of the arginine fingers of Rvb1 and Rvb2, we substituted these residues with alanine or lysine using sited directed mutagenesis and tested their impact on cell viability. Because *RVB1* and *RVB2* genes are essential for cell viability, we used a galactose-inducible/glucose-repressible promoter in *S. cerevisiae* to assess the effect of plasmid-borne *RVB1* or *RVB2*. Cells carrying plasmids were spotted on plates containing glucose, on which the endogenous gene expressions would be turned off. As it would be expected if the arginine finger played a crucial role, the expression of the *RVB1* arginine finger alanine variant (Rvb1-R366A) was lethal at both 30 °C and 25 °C. However, yeast cells expressing the *RVB2* arginine finger alanine variant (Rvb2-R350A) were viable at 30°C but showed significant growth defect at 25 °C, presenting a cold-sensitive phenotype (**Figure 2A**). While the growth defect of cells expressing Rvb2-R350A variant could be rescued at a higher temperature (37 °C), expression of Rvb1-R366A was not viable at any temperature. We, therefore, tested whether the variation of arginine fingers to a similarly charged amino acid, lysine, would be permissible and allow for growth. Interestingly, this analysis resulted in opposite growth phenotypes: while the expression of the lysine arginine finger variant of Rvb1 (Rvb1-R366K) slowed cell growth, the expression of the analogous variant of Rvb2 (Rvb2-R350K) was lethal (**Figure 2A**). The growth defect from the expression of both Rvb1 and Rvb2 R/K variants could be partially rescued at 37 °C.

**Figure 2.**
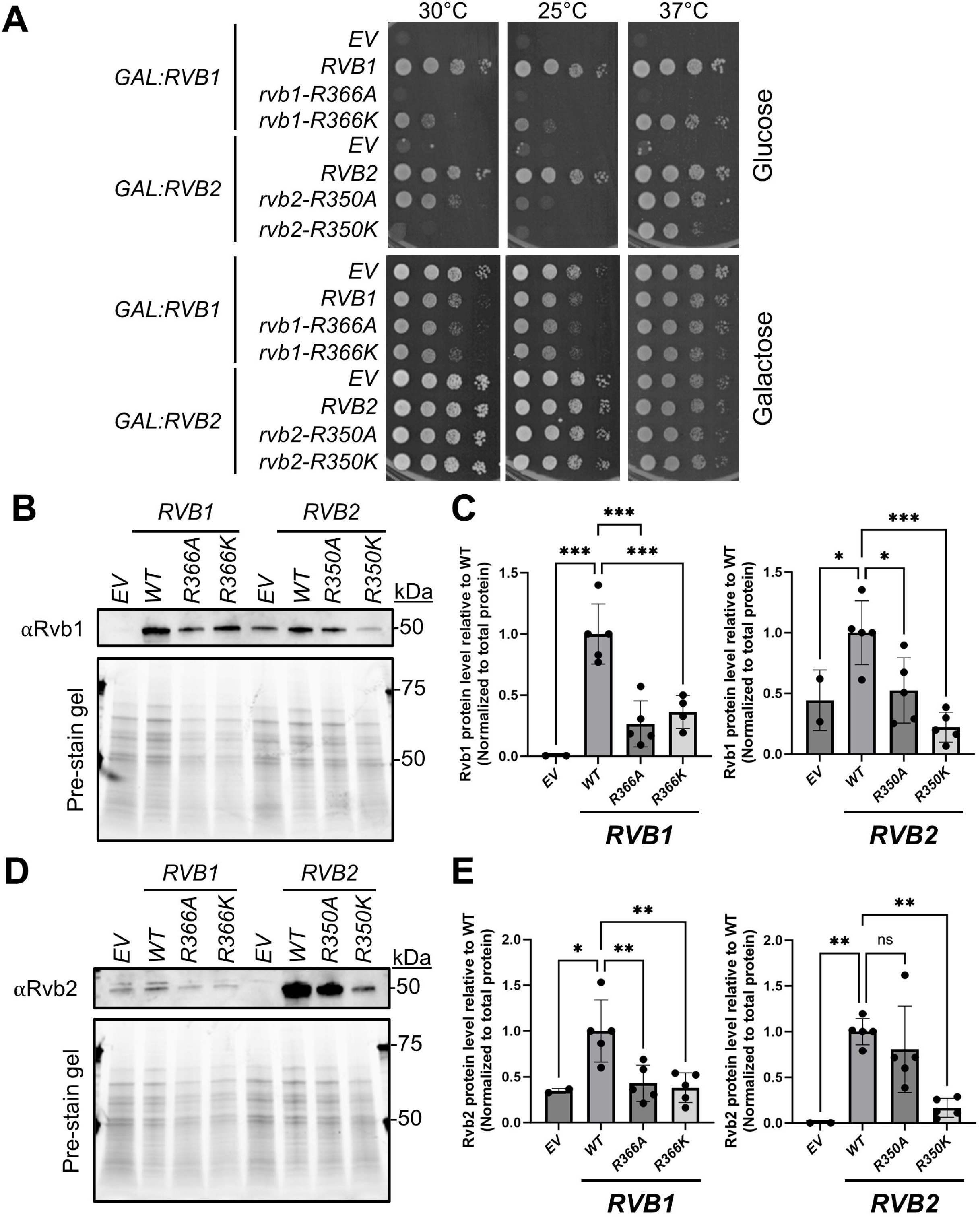
Arginine finger variants of the highly conserved Rvb1 and Rvb2 proteins differentially impact cell growth and steady-state protein levels. **A**) Cell growth was analyzed via serial dilutions, using a galactose-inducible/glucose-repressible promoter on *RVB1* or *RVB2* on media containing galactose (bottom panels) or glucose (top panels). **B)** Western blot analysis of steady-state Rvb1 levels in cells expressing EV or Rvb1/Rvb2 (as indicated) WT, or arginine finger RA or RK mutation. **C)** Quantification of **B**. Signal was normalized to total protein (using pre-stained gel) as a loading control and then normalized to cells expressing WT *RVB1* or *RVB2*, which was set to a value of 1. One-way ANOVA was then used to compare protein levels between the different variants. Error bars represent ± SD; ns, not significant; * *p* < 0.05; ** *p* < 0.01; *** *p* < 0.001; *N* = 5. **D)** Western blot analysis of steady-state Rvb2 levels in cells expressing EV or Rvb1/Rvb2 (as indicated) WT, or arginine finger RA or RK mutation. **E)** Quantification of **D**. Signal was normalized to total protein (using pre-stained gel) as a loading control, and then normalized to cells expressing WT *RVB1* or *RVB2*, which was set to a value of 1. One-way ANOVA was then used to compare protein levels between the different variants. Error bars represent ± SD; ns, not significant; * *p* < 0.05; ** *p* < 0.01; *** *p* < 0.001; *N* = 5.

These unexpected results highlighted the different impacts of the arginine fingers of Rvb1 and Rvb2 on cell growth, indicating their regulation or function may be distinct (**Figure 1C**). To corroborate this finding, we tested whether overexpression of *RVB1* and *RVB2* and their variants are differentially tolerated in wild-type yeast cells. Overexpression of both *RVB1* and *RVB2* and their arginine finger variants results in smaller colony sizes and varying extents of slower growth phenotype that is exacerbated at lower temperatures (**Figure S2A**). Interestingly, while overexpression of *RVB1* or *rvb1-R366K* causes a significant growth defect, this phenotype is not observed for cells overexpressing *rvb1-R366A*. When *rvb1-R366A* is the sole expressed variant of Rvb1 in cells, it does not support growth (**Figure 2A**). However, in cells expressing functional wildtype *RVB1*, the expression of *rvb1-R366A* does not hamper cell growth, suggesting Rvb1-R366A can’t compete with wildtype Rvb1 (**Figure S2A**). In addition, wildtype yeast cells tolerate the overexpression of *RVB2* better than *RVB1* (**Figure S2A**). Compared to wild-type *RVB2*, overexpression of the *rvb2-R350A* and *rvb2-R350K* variants caused slow growth at 30°C and 25°C. Overexpression of both R/K variants (*rvb1-R366K* and *rvb2-R350K*) significantly impacts growth in wildtype yeast, indicating the negative growth effects arising from arginine finger variants are related to the positive charge of the arginine finger variant. Collectively, these data suggest that Rvb1 and Rvb2 are differentially regulated in cells as their individual overexpression is differentially tolerated in cells and variations to their conserved arginine finger residues result in distinct cellular growth outcomes.

### Alanine variants of the arginine fingers in Rvb1 and Rvb2 destabilize the protein in vivo but not in vitro

To further understand the reason behind the different growth phenotypes arising from the expression of the Rvb1 and Rvb2 arginine finger variants, we measured the steady-state levels of Rvb1 and Rvb2 proteins in cells expressing wildtype or arginine finger variants (R/A and R/K). Because arginine finger protrudes from one protomer to the other, we reasoned that the respective variants may influence the complex stability by affecting intersubunit interactions between the protomers. Western blots revealed that expression of all arginine finger variants (Rvb1-R/A or R/K and Rvb2-R/A or R/K) reduced the steady-state levels of the Rvb1 protein (**Figure 2B-C**). Similarly, Rvb2 steady-state levels were decreased in cells expressing Rvb1 or Rvb2 R/K variants, but they did not change significantly in cells expressing the Rvb2-R350A variant (**Figure 2D-E**). Additional replicates of the representative blots shown in **Figure 2B** are provided in **Figure S2B**.

To assay the impact of arginine finger variants on protein folding and stability, we recombinantly expressed and purified the wildtype and variant Rvb1/2 complexes from *E. coli*. Wildtype Rvb1/2 complex or arginine finger variants (Rvb1/Rvb2-R350A, Rvb1-R366A/Rvb2, and Rvb1-R366A/Rvb2-R350A) were expressed and purified over three purification steps (affinity, ion exchange, and size exclusion chromatography) in identical manners (see methods). Several independent protein preparations were used for analytical assays to test homogeneity and stability. The chemical purity of purified complexes after the final step of purification was analyzed via Coomassie-staining on an SDS PAGE (**Figure S3A**). To assess the oligomerization status of purified Rvb1/2 and variants, we used dynamic light scattering (DLS). This analysis confirmed the results from size exclusion chromatography and showed a homogeneous population of purified Rvb1/2 complexes with similar radius for wildtype and variants (**Figure S3B**). To assess stability, we measured the thermal denaturation of purified and wild-type Rvb1/2 variants (**Figure S3C-D**). We did not observe any major changes between the apparent melting temperature of the isolated complexes (**Figure S3D**). However, there is a slight shift towards lower overall stability of the individual R/A variants compared to the wildtype and a change in the melting pattern of the double R/A variant compared to control (**Figure S3C**). Together, our data suggest that individual arginine finger variations can alter the steady-state levels of the proteins *in vivo* but have little impact on the complex stability *in vitro*.

### Alanine variants of the arginine fingers in Rvb1 and Rvb2 differentially impact the ATPase activity of the complex

To assess if arginine finger variants of Rvb1 and Rvb2 impact the enzyme’s activity, we characterized the wild-type and variant Rvb1/2 complexes *in vitro*. We first measured the steady-state ATPase activity of Rvb1/2 and its R/A variants using an NADH-coupled ATPase assay under multiple turnover conditions (56). Using the indicated ATP concentrations in these assays, we obtained *k*_cat_ and *K*_M_ values (**Figure S4A**). The ATPase activity of wildtype Rvb1/2 is low at 0.84 ± 0.02 molecules of ATP per minute, similar to what has been reported before (10, 18, 28, 37, 38, 57). Interestingly, however, the Rvb1 arginine finger alanine variant increases the complex’s ATPase activity, with the Rvb1-R366A/Rvb2 complex hydrolyzing 1.2± 0.04 molecules of ATP per minute. In contrast, complexes based on the Rvb2 arginine finger variant and the double alanine variant (Rvb1/Rvb2-R350A; Rvb1-R366A/Rvb2-R350A) show very low ATPase activity (**Figure 3A-B**). Single turnover assays, measuring phosphate hydrolysis over time after incubating excess ATPase concentrations with limiting amounts of ATP-γP^32^, show no significant difference in Pi hydrolysis by either of the single R/A variants (**Figure 3C-D**), indicating that the tested arginine fingers are not critical for ATP hydrolysis. Collectively, these data indicate that although highly similar, the arginine fingers of Rvb1 and Rvb2 have distinct contributions to the ATP hydrolysis of the complex.

**Figure 3.**
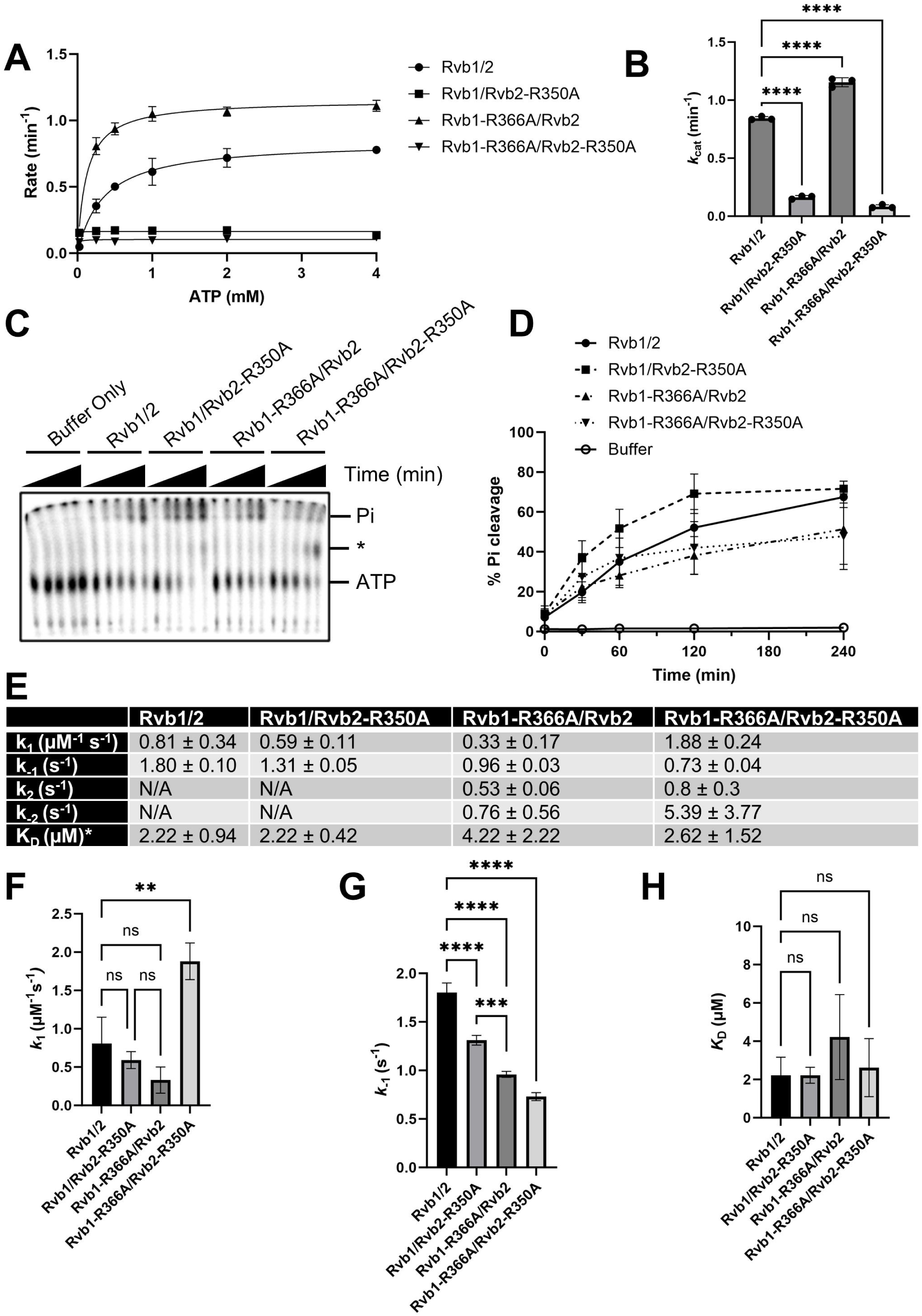
The arginine finger variants of Rvb1 and Rvb2 have different effects on ATP binding, hydrolysis, and release of the Rvb1/2 complex. **A**) Steady-state ATPase rate was measured via an NADH-coupled assay. Shown is the calculated ATPase rate plotted at different concentrations of ATP for each Rvb1/2 variant. A no-ATPase control was subtracted from the calculated rates as background. **B)** The *k*_cat_ calculated from the curve shown in **A** of ATP turnover over time at different ATP concentrations for the different Rvb1/2 variants. One-way ANOVA analysis was used to compare *k*_cat_ between purified Rvb1/2 and arginine finger mutants. Error bars represent ± SD; ns, not significant; **** *p* < 0.0001; *N* = 3. **C)** Single turnover kinetics of Rvb1/2 and arginine finger mutants was measured by analyzing the hydrolysis of ATP using radiolabeled ATP [γP^32^] over time. * Some radioactivity was detected to migrate between Pi and ATP on the TLC plate, which may be Pi-Pi. **D)** Cleavage of Pi was calculated by quantifying density of Pi (top band) divided by the density of the total lane from **C** and multiplying by 100 to obtain a % Pi cleavage. Five replicates, from independent protein preparations, were analyzed. **E)** Association and dissociation rate constants calculated for Rvb1/2 and variants. For Rvb1/2 and Rvb1/Rvb2-R350A, *K*_D_ was calculated as *K*_D_=*k*_-1_/*k*_1_. For Rvb1-R366A/Rvb2 and Rvb1-R366A/Rvb2-R350A, *K*_D_ was calculated after the fraction bound and *k*_-2_=*K*_D_\**k*_2_\**k*_1_/*k*_-1_. **F-G)** Association (**F**) and dissociation (**G**) rate constants for binding of Rvb1/2 and variants to mant-ATP. Error bars represent the error in fit of the data (**Figure S4B-C**) to a curve; ns, not significant; ** *p* < 0.01; *** *p* < 0.001; **** *p* < 0.0001. **H)** Comparison of dissociation constant (*K*_D_) for each Rvb1/2 variant with mant-ATP. One-way ANOVA was performed to compare the dissociation constants. Error bars represent error in the fit to the curve; ns, not significant.

### Arginine fingers of Rvb1 or Rvb2 are differentially regulated in the active sites of the complex

To understand how alanine substitutions of the arginine fingers within Rvb1 or Rvb2 affect the ATPase activity of the complex, we measured the rate constant for mant-ATP dissociation (*k*_off_) or mant-ATP association (*k*_on_) (**Figure 3E-G**, **Figure S4B-C**). We did not observe any significant change in the association of mant-ATP to Rvb1-R366A/Rvb2 or Rvb1/Rvb2-R350A variants compared to wildtype control (**Figure 3F**). However, the association rate of mant-ATP binding to Rvb1-R366A/Rvb2-R350A is higher by nearly 2-fold, and the rate constant for ATP dissociation (*k*_off_) is approximately 2-fold lower than that of wildtype control. Thus, the overall affinity (*K*_D_) of Rvb1-R366A/Rvb2-R350A variant for mant-ATP compared to wildtype remains the same (**Figure 3E**). All variants have a slower dissociation rate of mant-ATP compared to wildtype control (**Figure 3G, S4C**). However, the observed changes in the dissociation rates of mant-ATP did not lead to a significant change in the dissociation equilibrium constants (**Figure 3E, H**). While analyzing the association data, we noted that the time dependence of the fluorescence signal recorded in experiments with Rvb1-R366A/Rvb2 and Rvb1-R366A/Rvb2-R350A could not be fit with a one exponential equation (**Figure S4B**). These fluorescence time courses were best fit with a two-exponential equation. We attribute this apparent biphasic interaction of the variants with mant-ATP to a potential rearrangement of the fluorophore (the mant group) and not to a second step of the binding event. Rvb1-R366 is ∼3 Å away from Rvb2-R397 in the sensor II motif (**Figure S4E**) and to the nucleotide bound to the Rvb2’s ATP binding site. Substitution of Rvb1’s arginine finger to alanine (Rvb1-R366A) would allow the fluorophore more flexibility to move into the binding site, causing the observed second phase. This is not the case for the same substitution in Rvb2, as in this case the equivalent arginine in Rvb1’s sensor II motif (Rvb1-R413) is more distant from the binding site (∼5 Å). Thus, two association and dissociation rate constants were calculated for the proteins containing the Rvb1 variant (Rvb1-R366A/Rvb2 and Rvb1-R366A/Rvb2-R350A; **Figure 3E**), and these values were used to calculate the *K*_D_. These observed differences prompted us to test whether there was an overall difference in the distance between the arginine finger and the bound nucleotide in the available structures of yeast Rvb1/2 (20, 22, 27, 39, 53–55). Overall, both arginine fingers are >5 Å away from the bound nucleotide in published structures and the Rvb1’s arginine finger is more distant (∼1.5 x) from the bound nucleotide than the Rvb2’s arginine finger (**Figure S4F**). Collectively, these data reveal that the alanine substitutions of the arginine fingers of Rvb1 and Rvb2 proteins have different effects on the complex’s ATPase activity and the dissociation of the nucleotide from the complex. Although seemingly similar, the arginine fingers within highly homologous Rvb1 and Rvb2 do not appear to be in functionally symmetric active sites even though the ATP binding sites do not have significantly different affinities for ATP.

### Molecular dynamics simulations reveal the differential effects of alanine substitution of Rvb1 and Rvb2 arginine fingers on the insertion domain

Rvb1/2 hydrolyzes ATP at its catalytic cores within a ring to achieve binding or conformational changes, often far from the active sites, within different biological contexts. Despite available structural information on the Rvb1/2 complex, the complex dynamics remain unknown. To gain insights into Rvb1/2 dynamics and the long-range effects from alanine substitutions of the arginine fingers within the complex, we turned to molecular dynamics (MD) simulations. To compare the dynamic ensemble of the arginine finger variants with wildtype control, we performed three independent 100 ns MD simulation replicates for each complex. RMSD indicates that the simulations reached equilibrium after 10 ns (**Figure S5A**), and RMSF analysis shows higher dynamics in the variants compared to the wildtype complex (**Figure S5B-C**). To compare the dynamics of the variants with the wildtype complex, we analyzed mean internal Cα distances from MD simulations. In this analysis, the internal distances between all Cα carbons were measured for every frame of the trajectory and an average was taken (**Figure 4**). The analysis revealed that overall, all variants experienced increased Cα carbon distances compared to the wildtype. Comparing Rvb1-R366A/Rvb2 and the double variant with wild-type, each variant is more dynamic in each protomer, including in the insertion domain, compared to wild-type (**Figure 4B, C** *lower panels*). Surprisingly, however, comparing Rvb1/Rvb2-R350A and wildtype, in one of the Rvb1 chains, there is significantly reduced dynamics (**Figure 4A**), which also results in the higher dynamics apparent for the same protomer in the variant. In contrast, comparing Rvb1-R366A/Rvb2 and wildtype, overall, the variant is significantly more dynamic than wildtype control (**Figure 4D**).

**Figure 4.**
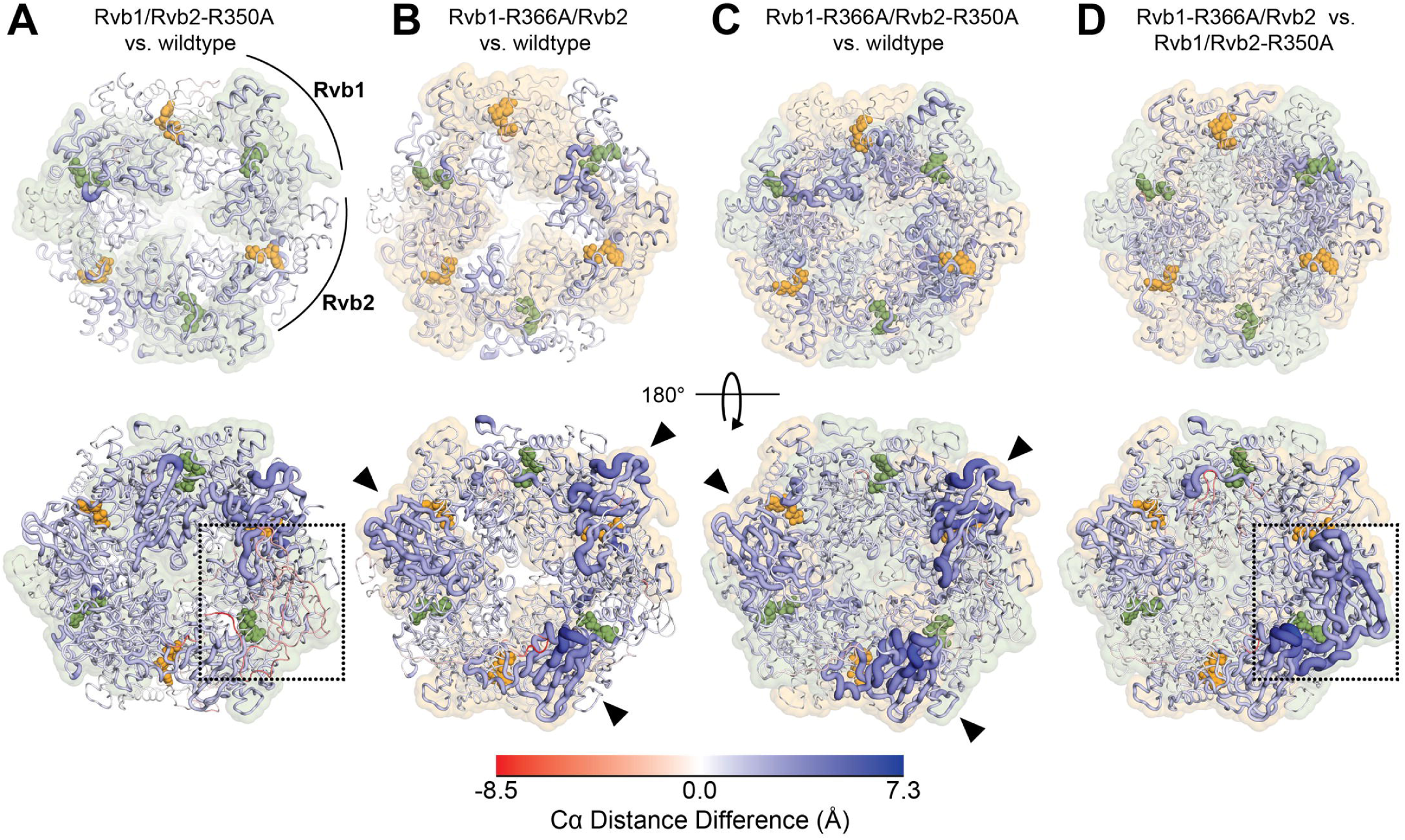
Mean internal Cα distance differences from MD simulations reveal the rigidity imposed by alanine substitution of the Rvb1’s arginine finger on the complex. Mean internal Cα distance differences between **A)** Rvb1/2 and Rvb1/Rvb2-R350A, **B)** Rvb1/2 and Rvb1-R366A/Rvb2, **C)** Rvb1/2 and Rvb1-R366A/Rvb2-R350A variant and **D)** Rvb1-R366A/Rvb2 and Rvb1/Rvb2-R350A from MD simulations plotted onto the last frame of the simulation. In all panels, ATP is shown as spheres in the active site. Putty thickness and color indicate distance differences where red indicates decreased distances in the mutant variant compared to wildtype and blue indicates increased distances in the variant compared to wildtype control. Boxes and arrows highlight measure changes.

Interestingly, the alanine substitution of the Rvb1 arginine finger increases the dynamics of the insertion domain of the protein (Domain II). While the variation is on Rvb1, all Rvb2 protomers become much more dynamic in a symmetric manner. Similarly, comparing Rvb1-R366A/Rvb2-R350A and wildtype, we observe symmetrically increased dynamics of Rvb2. To tease out the individual effects from alanine substitutions of arginine fingers in Rvb1 and Rvb2 proteins, we compared Cα carbon distances between Rvb1-R366A/Rvb2 and Rvb1/Rvb2-R350A. We again observe reduced dynamics of a single protomer of Rvb1 in the complex when Rvb1’s arginine finger is substituted with alanine. Overall, the alanine substitution of the arginine finger of Rvb2 causes asymmetric rigidity of the insertion domain of Rvb1 within the complex. This is observed in both Rvb1/Rvb2-R350A and Rvb1-R366A/Rvb2-R350A, both of which are catalytically inactive in steady-state conditions.

In summary, using MD simulations, we find that alanine substitutions of the arginine fingers near the active sites of Rvb1/2 in domain I of the protein have long-range impacts on dynamics of the domain II that is known to influence both the activity and interactions of the complex (7, 15, 17, 20, 22, 24, 25, 41, 49, 58, 59). These data strongly suggest that activity within the catalytic site of ATP hydrolysis is relayed to the insertion domain to influence the protein’s function. Further, the data points to the asymmetric functional active sites of the complex.

### Differential impact of the alanine substitution of arginine fingers of Rvb1 and Rvb2 on the function of the complex in snoRNP assembly

The insertion domain of Rvb1/2 is critical for binding of the complex to several proteins in different biological contexts (7, 11, 20, 22, 27, 39, 54, 60). Among the factors that interact with the insertion domain is the essential nucleolar protein Nop58, which is a core member of the box C/D snoRNP complexes (47). Within the R2TP complex (61), Tah1 and Pih1 serve as adapters for binding of the complex to Nop58. The structural and biochemical studies of the complex of R2TP with Nop58 have revealed that Nop58 has direct interactions with Rvb1/2’s insertion domain (40, 41, 58, 62). To validate our MD simulation results showing an impact from alanine substitutions of Rvb1/2 arginine fingers on the insertion domain, we therefore tested how the arginine finger alanine substitution variants within Rvb1 and Rvb2 influence the stability of the Nop58 protein. Both arginine finger alanine substitution variants in Rvb1 and in Rvb2 reduce the overall steady-state levels of Nop58 (**Figure 5A-C**). In line with this, both variants result in a significant reduction of the steady-state levels of C/D snoRNAs (**Figure S6A-B**). A targeted high copy suppressor screen confirmed that the cold-sensitive phenotype of yeast cells expressing *rvb2-R350A*, can be partially rescued by overexpression on Nop58 (**Figure 5D**). While overexpression of *NOP58* causes a significant slow growth phenotype in wildtype cells, *NOP58* overexpression in *rvb2-R350A* cells partially suppresses the growth defect of cells. Among snoRNP core factors, Nop58 is the only factor whose overexpression can suppress the cold-sensitive phenotype of *rvb2-r350A* cells (**Figure 5D**). The severe growth defect observed in cells expressing *rvb1-r366A* could not be rescued by overexpression of Nop58. In summary, these data indicate that the alanine substitution of the arginine fingers of Rvb1/2 impact snoRNP assembly by decreasing the steady-state levels of Nop58.

**Figure 5.**
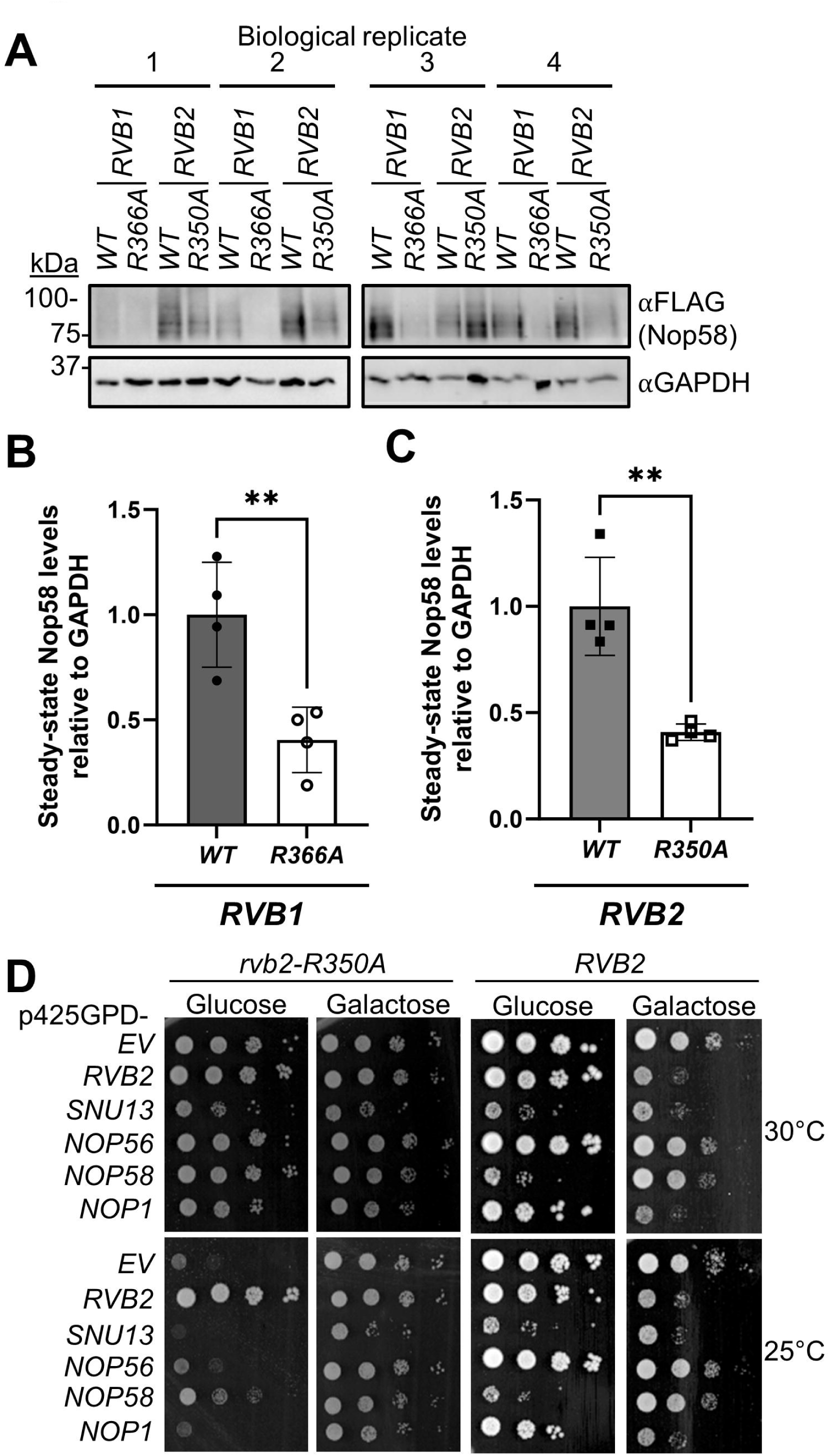
The arginine finger variants of Rvb1 and Rvb2 both affect the snoRNP biogenesis pathway. **A**) Western blot analysis of cellular levels of Nop58 (in *NOP58-5xFLAG* cells) in WT and arginine finger mutant cells. **B-C)** Quantification of Nop58 from blots shown in **A**. Signal was normalized to GAPDH loading control, and then normalized to cells expressing WT *RVB1* or *RVB2*, which was set to a value of 1. Unpaired t-test were then used to compare protein levels between *RVB1* and *rvb1-R366A* (**B**), and between *RVB2* and *rvb2-R350A* (**C**) expressing cells. Error bars represent ± SD; ns, not significant; * *p* < 0.05; ** *p* < 0.01; *N* = 4. **D)** Serial dilutions of *RVB2* or *rvb2-R350A* cells overexpressing snoRNP core proteins. While overexpression of *NOP58* leads to sick growth in WT cells, it rescues the cold sensitive phenotype observed in *rvb2-R350A* cells.

### Differential sensitivity of Rvb1 and Rvb2 to stressors

Rvb1/2 are residents and regulators of both mammalian and yeast stress granules (62–66). The role of Rvb1/2 in stress granules has been linked to gene expression regulation (64) as well as chaperone activity of Rvb1/2 (63, 65, 67). Using the known stressors that impact stress granules, we tested how cells expressing wildtype or the arginine finger variants of Rvb1/2 respond to cellular stress (**Figure 6**). We employed two different stress conditions that induce stress granule formation: glucose starvation-induced by 2-deoxy-D-glucose (2-DG) (63, 64, 67) and sodium azide (NaN_3_) treatment (63, 68–70). Because expression of Rvb1/R/A variant was not viable and combining that with stressors did not support cell growth, we included the R/K variants in this analysis. Our analysis shows that the yeast cells expressing *rvb1-R366K* variant become resistant to glucose starvation stress and grow about as well as the wildtype control cells (**Figure 6**). However, cells expressing *rvb2-R350A* are sensitive to glucose starvation. In contrast, we observe that *rvb2-R350A* cells are resistant to NaN_3_ treatment and grow comparable to wildtype control cells under this condition. While the *rvb1-R366K* cells are slightly less sick than control cells, they still grow slower than control cells on NaN_3_ media (**Figure 6**). The difference in stress response by Rvb1 and Rvb2 variants highlights the differential impact of these proteins on their activity within stress response pathways.

**Figure 6.**
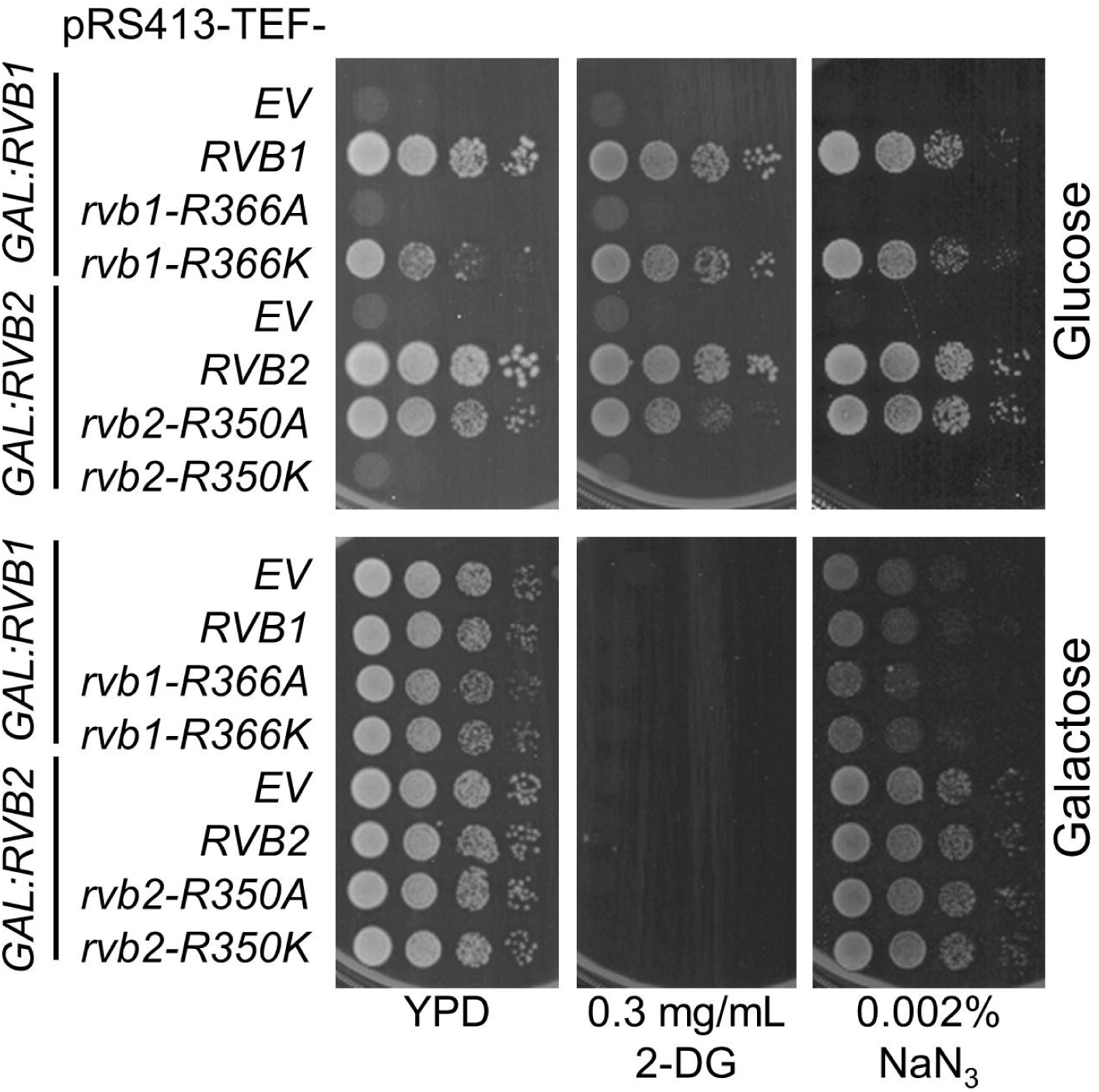
Differential sensitivity of Rvb1 and Rvb2 arginine finger mutants to stressors. Serial dilutions analyzing the response of Rvb1 and Rvb2 and their variants to glucose starvation via addition of 2-deoxy-D-glucose (2-DG) and sodium azide (NaN_3_) stress.

## Discussion

AAA^+^ ATPases couple binding, ATP hydrolysis, and phosphate release with conformational changes that allow for remodeling or chaperone activity (1–5, 71). Though AAA^+^ ATPases are a broad classification of proteins that are involved in different pathways, they all have conserved structural elements for catalysis (3, 4). The coordinated action of the Walker A, Walker B, sensor motifs, and arginine fingers within the multi-subunit ATPase ring is important for the function of the complex. Among these features, arginine fingers are thought to be directly responsible for “starting” the AAA motor and to be involved in the crosstalk between individual protomers of the complex (6, 72–76). There are different proposed mechanisms for the coordination of ATP hydrolysis, including synchronized, rotational, and sequential mechanisms (71). Regardless of the mode, the hydrolysis coordination between the subunits is likely sensed primarily by the arginine fingers (74). Much evidence also suggests that the arginine fingers are essential in controlling the directionality of AAA motors that have a rotating or revolving mechanism of action (74–77). Thus, variations to the arginine fingers of AAA ATPases are expected to disrupt the complex function and ATPase activity.

Here, we provide evidence that Rvb1 and Rvb2, two closely related and highly homologous proteins that work together as part of the Rvb1/2 AAA^+^ ATPase complex, have different regulatory functions in the cellular context. Our data reveal that variations to the arginine fingers of Rvb1 and Rvb2 have specific effects both *in vivo* and *in vitro*. The arginine fingers of Rvb1/2, while also contributing to the ATP related enzymatic properties of the complex (**Figure 3**), could also be involved in assembly of Rvb1/2 hexamers. There is some evidence that arginine fingers drive formation of ATPase dimers and promote intersubunit interaction during assembly (74). Despite a large overlapping interface, Rvb1/2 dimers may need to assemble via key interactions of their arginine fingers, and thus promote hexameric complex formation. Upon substitution of the arginine with other residues, hexameric formation could be compromised. This could explain why we observe reduced levels in both Rvb1 and Rvb2. It is possible that monomeric Rvb1 and Rvb2 proteins are degraded in vivo if not assembled into oligomers. To date, there is no evidence that monomeric Rvb1 or Rvb2 have a biological function. Another possibility is that the turnover of the complex is triggered due to disruption of its interaction with other interactors.

Our results hint toward the advantages of protomer diversification in AAA^+^ ATPases. While some AAA^+^ ATPases, such as Cdc48 and RuvB, form homomeric rings (42, 73, 78), many including Rvb1/2 and the AAA^+^ ATPase ring of the 26S proteasome form heteromeric rings (79, 80). Why some AAA+ ATPases form heteromeric rings, and others do not is largely unknown. In the case of Rvb1/2, the individual protomer ATPase activity rates are significantly different (10) and the Rvb1 homohexamer has been shown to be catalytically inactive (12). In line with this, our data indicate that the Rvb1’s arginine finger is insufficient to activate the complex and *in vitro*, in the absence of any other factors, the Rvb1 protomers rely on the Rvb2’s arginine finger for ATPase activity (**Figure 3**). In contrast, deactivation of Rvb1’s arginine finger is tolerated by the complex and does not hamper the complex ATPase activity *in vitro*. While the arginine finger variations have no impact on the binding of the nucleotide, both variations affect the *k*_off_ of ADP release (**Figure 3**). These data strongly suggest that analogous arginine fingers in Rvb1 and Rvb2 contribute to regulation of the catalytic activity of the complex unequivalently.

Based on structural data, it is likely that binding of a cofactor could induce conformational changes in Rvb1/2 that could push the arginine finger into the active site. In fact, many published structures in which Rvb1/2 is bound to a cofactor or complex show shorter distances between arginine fingers and bound nucleotide (6-9Å) (20, 22, 27, 39, 53–55) as compared to structures where Rvb1/2 is not bound to a cofactor or in a larger complex (12-16Å) (16, 17, 49, 51, 52). This mechanism of pushing the arginine finger into the active site is similar to how GTPases are activated by GTPase-activating proteins (GAPs) (8, 9, 81, 82). GAPs have conserved arginine residues that are inserted into GTP-binding domains of GTPases that induce nucleotide hydrolysis. This “arginine finger hypothesis” (81) observed in GTPases could be a conserved regulatory mechanism for some ATPases. Such a mechanism for ATP hydrolysis could act as a means of regulation for the downstream effects of Rvb1/2. It has been shown that binding of cofactors to the insertion domain affects the conformation of the ATP binding domain, typically triggering release of nucleotide from the Rvb2 protomers (26, 48, 49). This in turn impacts the ATPase activity of Rvb1/2. In the case of INO80, the published structures show a shorter distance from the arginine fingers to bound nucleotide (∼6Å) (20, 22, 39, 53), and also show an increase in ATPase activity (37). On the other hand, binding of DHX34 shows an increase in distance between the arginine finger and bound nucleotide (∼13Å) and a decrease in ATPase activity (48). Additionally, it has been shown that the nucleotide-bound state of Rvb1/2 in turn affects the conformation of the insertion domain (7, 15, 17). This conformational change could alter the affinity of Rvb1/2 to cofactors. This allosteric communication between the ATP binding domain and the insertion domain is also confirmed by our MD simulation data (**Figure 4**). For example, ATP hydrolysis of Rvb1/2 could trigger the release of the mature snoRNP complex by leading to a conformational change in the Rvb1/2 insertion domains that has decreased affinity for bound snoRNPs. This could explain why elevated activity (such as with Rvb1-R366A variant; **Figure 3**) causes lethality *in vivo* (**Figure 2**) as such elevated ATPase activity could lead to release of immature complexes. We observe a drastic difference in the symmetry of the Rvb1/2 complex dynamics depending on whether the Rvb1 or Rvb2 arginine finger is mutated, suggesting alterations to the overall complex assembly (**Figure 4**). This asymmetry could also explain why mutation of the arginine finger in Rvb1 is lethal while the same mutation in Rvb2 is viable yet catalytically inactive and renders the cells cold-sensitive (**Figure 2**).

While Rvb1 and Rvb2 work together in a complex, they interact with cofactors in different manners. Immunopurification experiments pulling down individual proteins have shown that while Rvb1 and Rvb2 interact with many of the same proteins, some proteins are only pulled down by either Rvb1 or Rvb2 (83). Even for proteins that interact with both Rvb1 and Rvb2, the specific interactions with individual protomers are distinct. For example, the N-terminal domain of Nop58 mainly interacts with Rvb1 while its C-terminal domain interacts more with Rvb2 (41). Many other structural studies have shown that certain cofactors bind to the Rvb1/2 hexamer in an asymmetric manner (20, 26, 37, 41, 49). This asymmetric binding could result in different changes to protomer conformation, alteration to ATP binding or hydrolysis, or in protein flexibility or rigidity between Rvb1 and Rvb2 depending on which protomers are interacting with another protein. For example, binding of PIH1D1 to a RUVBL2 protomer in a hexameric ring causes conformational changes that promote dissociation of nucleotide and has been postulated to increase nucleotide exchange (49). Any differences between Rvb1 and Rvb2 could allow for differential regulation of Rvb1 and Rvb2 subunits in complex providing further modes of regulation.

Under the stress conditions we tested, we observed a difference in growth between Rvb1 and Rvb2 mutants (**Figure 6**). These interesting data suggest that Rvb1 and Rvb2 have different roles in stress response or are regulated differently in the cell in response to different environmental conditions. Such differences could have the potential to be exploited in certain disease models for therapeutic purposes. For example, RUVBL ATPases have been shown to have chaperone activity involved in disassembly of large aggregates in neurodegenerative disease models (65). They were previously reported to have disaggregase behavior, and downregulation of RUVBL1 using siRNA knockdown was shown to increase toxicity of aggregates (66). Many AAA ATPases have been identified as both biomarkers and potential therapeutic targets (84, 85). In fact, there have been a few small-molecule inhibitors identified for RUVBL1/2 with certain specificities (86–91). Further understanding of the differences between the activities and molecular regulation of these conserved ATPases could, therefore, inspire new modes of drug design to specifically modulate the complex’s activity within different contexts.

## Author Contributions

J.L.W and H.G designed the research; J.L.W, J.A.B, S.M.N, M.H, A.V, H.J.W, S.K. and H.G performed research and analyzed data; J.L.W, J.A.B, S.M.N, M.H, and A.V. contributed to writing sections of the first manuscript’s draft. J.L.W and H.G wrote the manuscript. All authors contributed to the final revision of the manuscript.

## Competing Interest Statement

The authors declare no competing interest.

## Classification

Biological Sciences, Biochemistry

## Acknowledgments

This work was supported by NIH Grant 1R35GM138123 (to H.G.). S.M.N was supported by an IRACDA fellowship (NIH 5K12GM000680-24). We thank Dr. Walid Houry at the University of Toronto for the generous gift of Rvb1 and Rvb2 antibodies. The wildtype Rvb1/2 bacterial expression plasmid was kindly provided by Dr. Karl-Peter Hopfner. We thank Drs. Elizabeth Draganova and Bo Liang (Emory, Dept of Biochemistry) for sharing their DLS equipment and expertise on DLS measurements and data analysis. We thank Sarah Webster and Zachary Bressman for their comments on the manuscript. This work used jetstream2 at Indiana University through allocation BIO240045 from the Advanced Cyberinfrastructure Coordination Ecosystem: Services & Support (ACCESS) program, which is supported by National Science Foundation grants #2138259, #2138286, #2138307, #2137603, and #2138296.

## Notes

https://github.com/hghalei/Rvb1-2

### Competing Interest Statement

The authors have declared no competing interest.

