## Supporting Information for "Differential roles of putative arginine fingers of AAA^+^ ATPases Rvb1 and Rvb2"

**This file includes:**

Supplementary Tables S1 to S3
Figure legends S1 to S6
Supplementary Figures S1 to S6

**Table S1. List of yeast strains used in this study.**

| Strain Name | Genotype | Reference |
|---|---|---|
| BY4741 | MATa; ura3Δ0; leu2Δ0; his3Δ1; met15Δ0 | |
| YHG010 | *BY4741; GAL1:RVB2:HYG* | This study |
| YHG495 | *BY4741; GAL:RVB1:KAN* | This study |
| YHG676 | *BY4741; NOP58-5xFLAG:NAT* | This study |
| YHG704 | *BY4741;NOP58-5xFLAG:NAT;GAL:RVB2:HYG* | This study |
| YHG706 | *BY4741; NOP58-5xFLAG:NAT; GAL:RVB1:KAN* | This study |

**Table S2. List of plasmids used in this study.**

| Plasmid Name | Backbone | Gene | Purpose |
|---|---|---|---|
| HG131 | pET28a | His-TEV-Rvb1,Rvb2 | Purification |
| HG176 | pET28a | His-TEV-Rvb1,Rvb2-R350A | Purification |
| HG184 | pET28a | His-TEV-Rvb1-R366A,Rvb2-R350A | Purification |
| HG185 | pET28a | His-TEV-Rvb1-R366A,Rvb2 | Purification |
| HG301 | pRS413TEF | Rvb2 | Overexpression |
| HG1016 | pET28a | His-TEV-Rvb1-R366K,Rvb2 | Purification |
| HG1017 | pET28a | His-TEV-Rvb1,Rvb2-R350K | Purification |
| HG3184 | pRS425GPD | Nop56 | Overexpression |
| HG3186 | pRS425GPD | Snu13 | Overexpression |
| HG3188 | pRS425GPD | Nop1 | Overexpression |
| HG3207 | pRS425GPD | Rvb2 | Overexpression |
| HG3250 | pRS415TEF-3xHA | Bcd1 | Immunopurification |
| HG3253 | pRS413TEF | Rvb2-R350A | Overexpression |
| HG3257 | pRS413TEF | Rvb1 | Overexpression |
| HG3258 | pRS413TEF | Rvb1-R366A | Overexpression |
| HG3313 | pRS425GPD | Nop58 | Overexpression |
| HG3315 | pRS425GPD | Rvb1 | Overexpression |
| HG3319 | pRS413TEF | Rvb1-R366K | Overexpression |
| HG3320 | pRS413TEF | Rvb2-R350K | Overexpression |

**Table S3. List of primers used in this study.**

| Primer | Sequence | Purpose |
|---|---|---|
| U14 | CGATGGGTTCGTACTCCTACCGTGG | Northern Blotting |
| U24 | GGTATGTCTCATTCGGAACTCAAAGTTCCATCTGAAGTAGC | Northern Blotting |
| snR8 | CACTCGCGCAGCTACCGATCTGGGCCAATGGGAGAC | Northern Blotting |
| snR35 | GAACAAAATGATGATCTCTCCGATGGACTTGACGC | Northern Blotting |
| snR46 | CTTCCCTTTGGAAATCGGAAATTCAATGATATGCCCTATGCCC | Northern Blotting |
| snR48 | GGAGAGTACTTAAACTTCACATCCTAACATTAGAGATGCCAG | Northern Blotting |
| snR51 | TGTAGTCATCAATTAGCCCC | Northern Blotting |

| | | |
|---|---|---|
| MRP | AATAGAGGTACCAGGTCAAGAAGC | Northern Blotting |
| Rvb2-350A | CCTCTCGATCTTTTGGATGCGTCAATTATTATTACAAC | Site-directed mutagenesis |
| RVB1-366A | CCACCTGATTTGATCGATGCATTGTTAATTGTTCGTAC | Site-directed mutagenesis |
| Rvb1-366K | GTGCCACCTGATTTGATCGATAAATTGTTAATTGTTCG TACATTAC | Site-directed mutagenesis |
| Rvb2-350K | GTTACCTCTCGATCTTTTGGATAAGTCAATTATTATTA CAACTAAAAG | Site-directed mutagenesis |

**Figure S1. Structural conservation of yeast Rvb1 and Rvb2 and human RUVBL1 and RUVBL2.** Sequence conservation map of yeast Rvb1 and Rvb2 and human RUVBL1 and RUVBL2 showing the conservation status of the sequences between *S. cerevisiae*, *H. sapiens*, and other eukaryotes. Conservation scale indicating level of sequence conservation shown below. Figure made using Consurf. Key ATPase sequence motifs indicated by colored lines or arrows.

**Figure S2. Although highly conserved, mutation to the arginine finger of Rvb1 and Rvb2 have different impacts on cell growth and protein level. A)** Serial dilutions of WT cells overexpressing *RVB1*, *RVB2*, or the arginine finger mutants. **B)** Biological replicates for quantifications shown in **Figures 2C,E**. Steady-state protein levels of Rvb1 and Rvb2 in *GAL:RVB1* and *GAL:RVB2* cells expressing EV, WT, or arginine finger mutants RA or RK was analyzed via western blot analysis.

**Figure S3. The arginine finger variants of Rvb1 and Rvb2 do not impact *in vitro* protein stability, conformation or homogeneity. A)** The purity of the purified Rvb1/2 hexamer (and variants) was compared on a Coomassie-stained SDS PAGE gel. A total of 2 µg of purified protein was loaded onto an SDS PAGE gel, electrophoresed until the dye front ran off, and then the gel was stained for protein with GelCode blue (Coomassie). **B)** To assure homogeneity in oligomerization of purified Rvb1/2 and variants, each purified hexamer was analyzed via dynamic light scattering (DLS). The radius of purified Rvb1/2 and variants (at indicated concentrations) was measured by dynamic light scattering using a Dynapro Plate Reader III. $N$ = 12. **C)** The thermal stability of purified Rvb1/2 variants was measured via the inflection temperature using a Tycho nanotemper instrument. Shown is the first derivative of the ratio of absorbance at 350 nm to the absorbance at 330 nm over increasing temperatures. The first derivative of the 350/330 nm ratio was normalized to the value at the calculated melting temperature, that value being set to 100%. **D)** Comparison of the average inflection temperature and standard deviation as measured in **C** for two different purified batches of each Rvb1/2 variant ($N$ = 2).

**Figure S4. Mutation of the arginine finger of Rvb1 and Rvb2 have different effects on ATP binding, hydrolysis, and release. A)** The $k_{cat}$ and $K_M$ values calculated for purified Rvb1/2 and arginine finger mutants via NADH coupled assay (**Figure 3A-B**). Shown are the mean values and their standard deviation. $N = 3$; n. c. not calculated due to insufficient curve. **B)** Association curves of mant-ATP with Rvb1/2 and variants, titrating in increasing concentrations of mant-ATP (black arrow indicating increasing concentrations of mant-ATP). **C)** Dissociation curves for Rvb1/2 and variants with mant-ADP. **D)** Structure of ATP binding pockets of Rvb2 (top panel) and Rvb1 (bottom panel) (PDB 6GEJ). Mutated arginine finger shown on the very left. Nearby arginine finger referred to in the text shown in the center. Individual subunits colored as in **Figure 1**. ADP molecule shown in grey and Mg ion shown in black. Dashed lines indicate measurements between residues and ligands. **E)** Average distance from arginine finger to bound ADP in published structures. PDBs: 5OAF, 6GEJ, 6GEN, 6FHS, 6FML, 7ZI4, 8ETS, 8ETU, 8ETW, 8EU9, 8EUF, 8OOC, 8OOR, 8OOP, 8X19, 8X1C. Unpaired t-test was then used to compare the average distance (of Rvb1 or Rvb2 protomers in one structure) between guanidine group of the arginine finger and the beta phosphate of bound nucleotide. * $p < 0.05$; $N = 16$.

**Figure S5. Root-Mean-Square Deviation (RMSD) and Root-Mean-Square Fluctuation (RMSF) calculations from MD simulations. A)** Root-mean-square deviation (RMSD) for wildtype, Rvb1-R366A/Rvb2, Rvb1/Rvb2-R350A, and Rvb1-R366A/Rvb2-R350A calculated from 100 ns molecular dynamics simulations. All simulations reached equilibrium at ~10 ns. **B)** Root-mean-square fluctuations (RMSFs) of Cα atoms for wildtype and the alanine substitution variants. **C)** RMSF difference comparing the alanine substitution variants to wildtype. The shaded area is 2 standard deviations above or below the average. Values above x = 0 indicate the mutant is more flexible than wildtype while values lesser than x = 0 indicate the mutant is less flexible than wildtype.

**Figure S6. The arginine finger variants of Rvb1 and Rvb2 both affect the snoRNP biogenesis pathway. A)** Northern blot analysis of box C/D and box H/ACA snoRNAs in cells expressing WT or arginine finger mutant Rvb1 or Rvb2. **B)** Quantification of snoRNA levels shown in (**A**). Signal was first normalized MRP loading control, and then normalized to cells expressing WT *RVB1* or *RVB2,* which was set to a value of 1. Two-way ANOVA was then used to compare protein levels between *EV*, *RVB1* and *rvb1-R366A*, and between *EV*, *RVB2* and *rvb2-R350A* expressing cells. Error bars represent ± SD; $N = 4$.

#### Figure S1

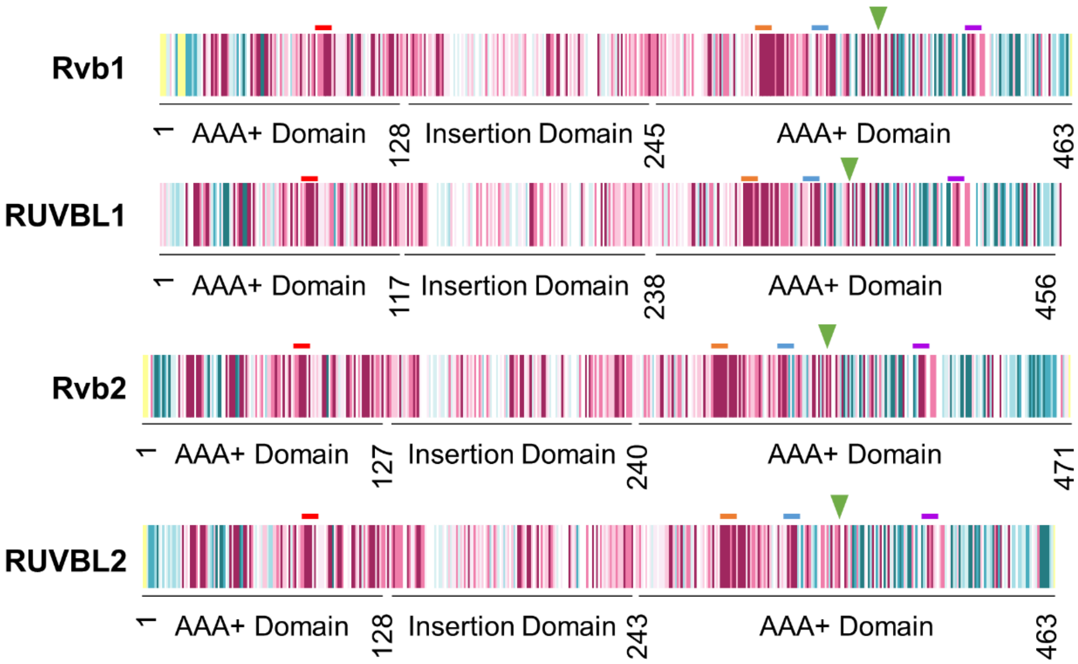


| | Rvb1 | RUVBL1 | Rvb2 | RUVBL2 |
|---|---|---|---|---|
| Walker A | 79-86 | 70-77 | 75-82 | 77-84 |
| Walker B | 306-312 | 298-303 | 292-297 | 295-300 |
| Sensor I | 337-343 | 327-334 | 322-328 | 326-330 |
| Arginine finger | 366 | 357 | 350 | 353 |
| Sensor II | 409-415 | 401-405 | 394-398 | 397-400 |

**The conservation scale:**

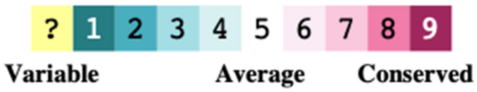

### Figure S2

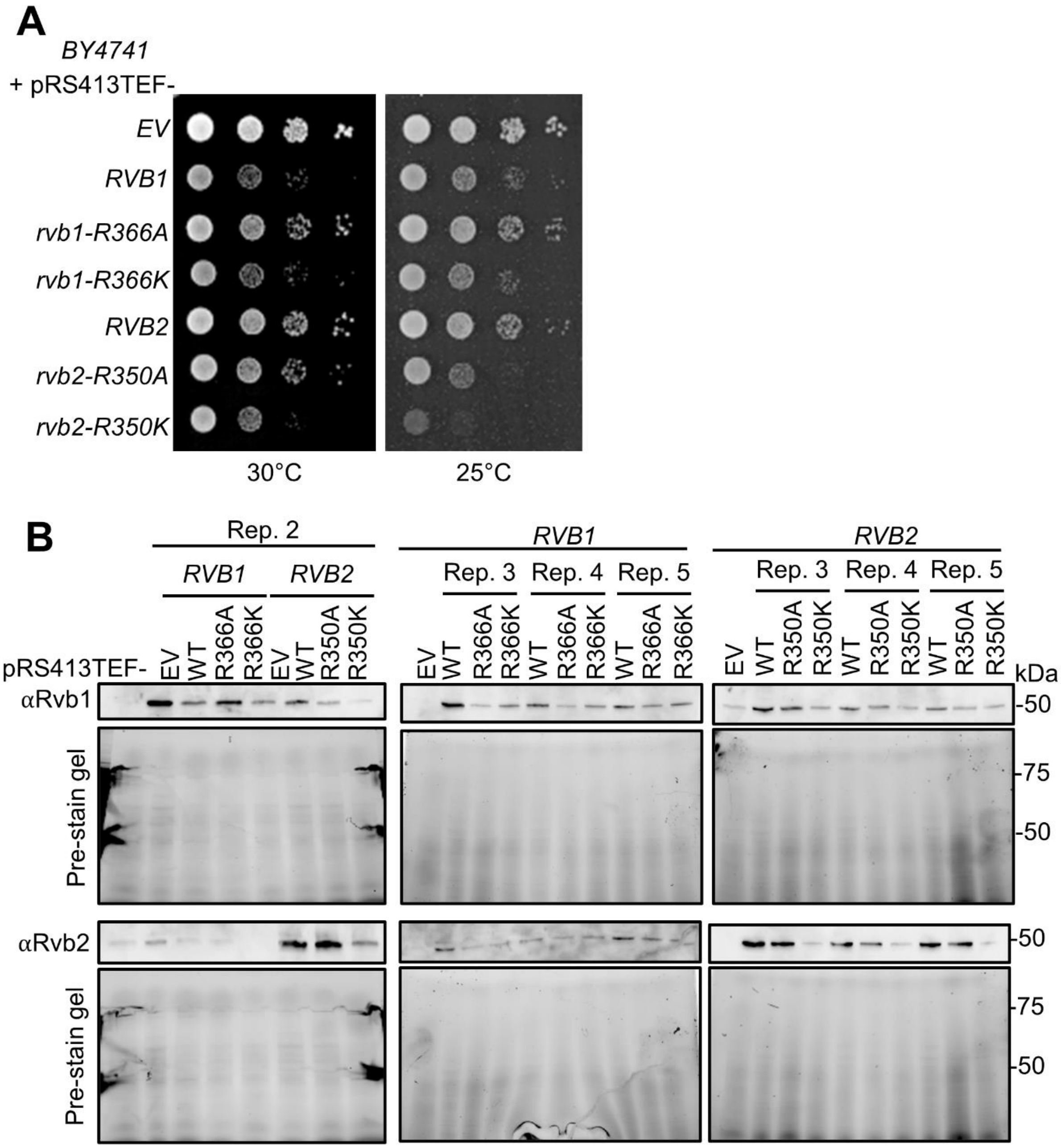
A
BY4741
+ pRS413TEF-
EV
RVB1
rvb1-R366A
rvb1-R366K
RVB2
rvb2-R350A
rvb2-R350K
30°C
25°C
B
Rep. 2
RVB1
RVB2
Rep. 3
Rep. 4
Rep. 5
pRS413TEF-
EV
WT
R366A
R366K
R350A
R350K
kDa
αRvb1
αRvb2
Pre-stain gel
-50
-75

### Figure S3

A

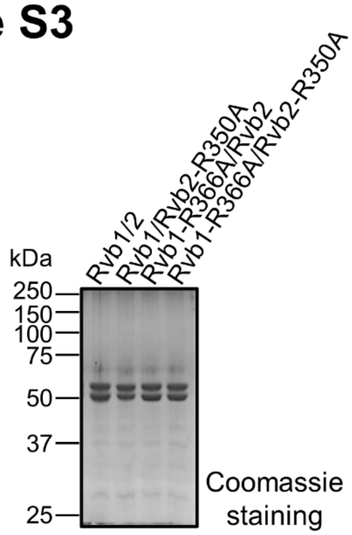


B

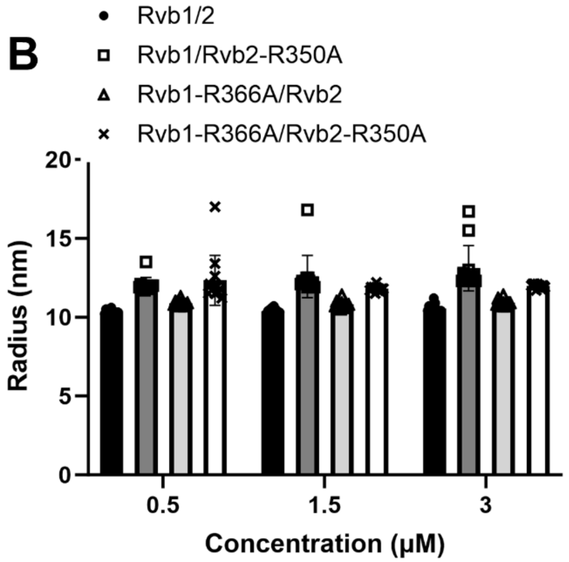


C

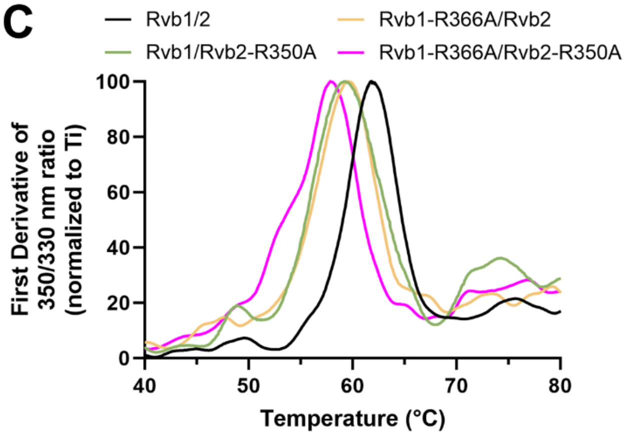


D

| Protein | Tm (°C) |
|---|---|
| Rvb1/2 | 61.7 ± 0.42 |
| Rvb1/Rvb2-R350A | 60.8 ± 1.91 |
| Rvb1-R366A/Rvb2 | 59.9 ± 0.42 |
| Rvb1-R366A/Rvb2-R350A | 57.9 ± 0.07 |

### Figure S4

A

| | $k_{cat}$ (min$^{-1}$) | $K_M$ (mM) |
|---|---|---|
| **Rvb1/2** | 0.84 ± 0.02 | 0.35 ± 0.02 |
| **Rvb1-R366A/Rvb2** | 1.15 ± 0.04 | 0.12 ± 0.01 |
| **Rvb1/Rvb2-R350A** | 0.16 ± 0.01 | n. c. |
| **Rvb1-R366A/Rvb2-R350A** | 0.08 ± 0.02 | n. c. |

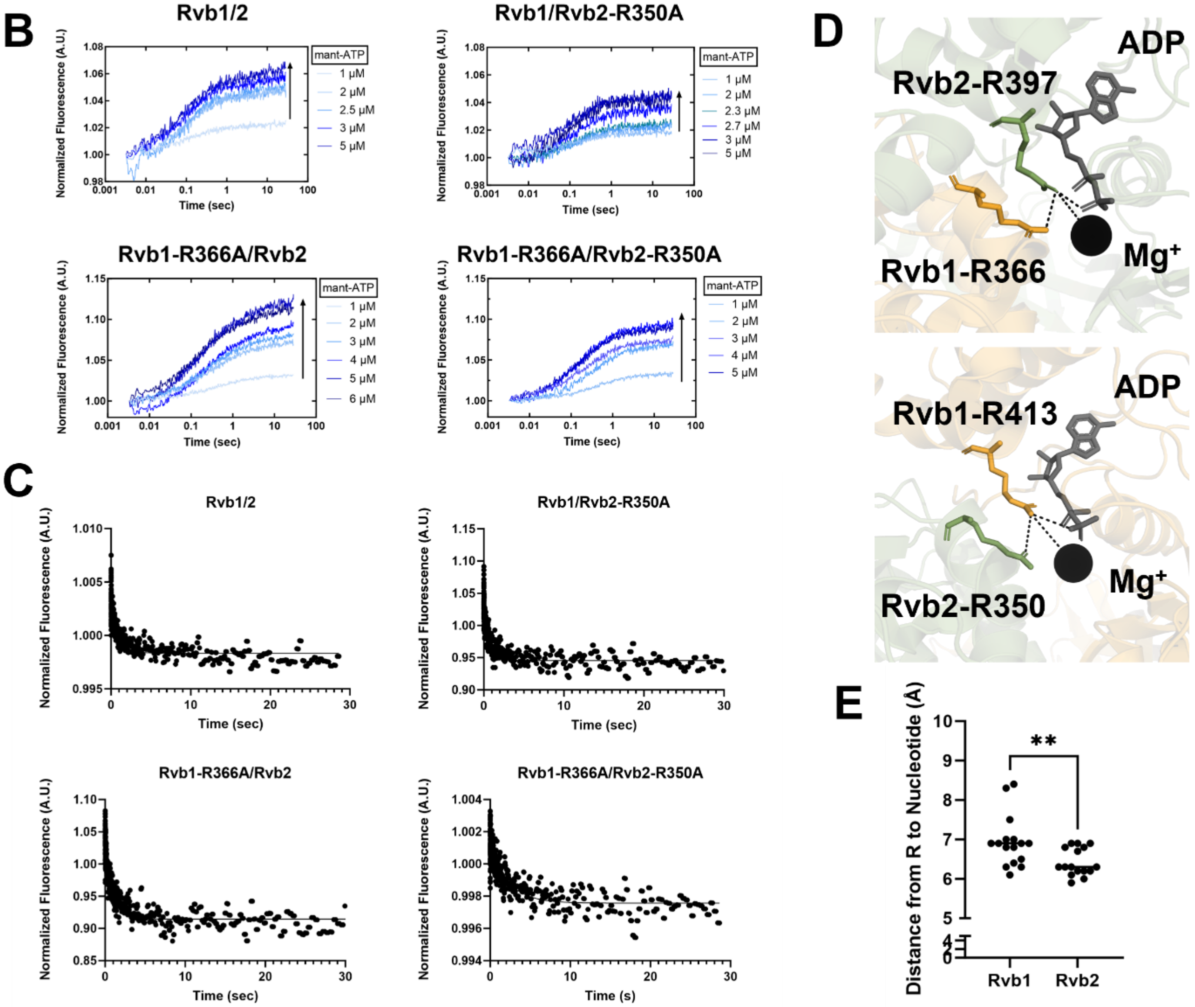

### Figure S5

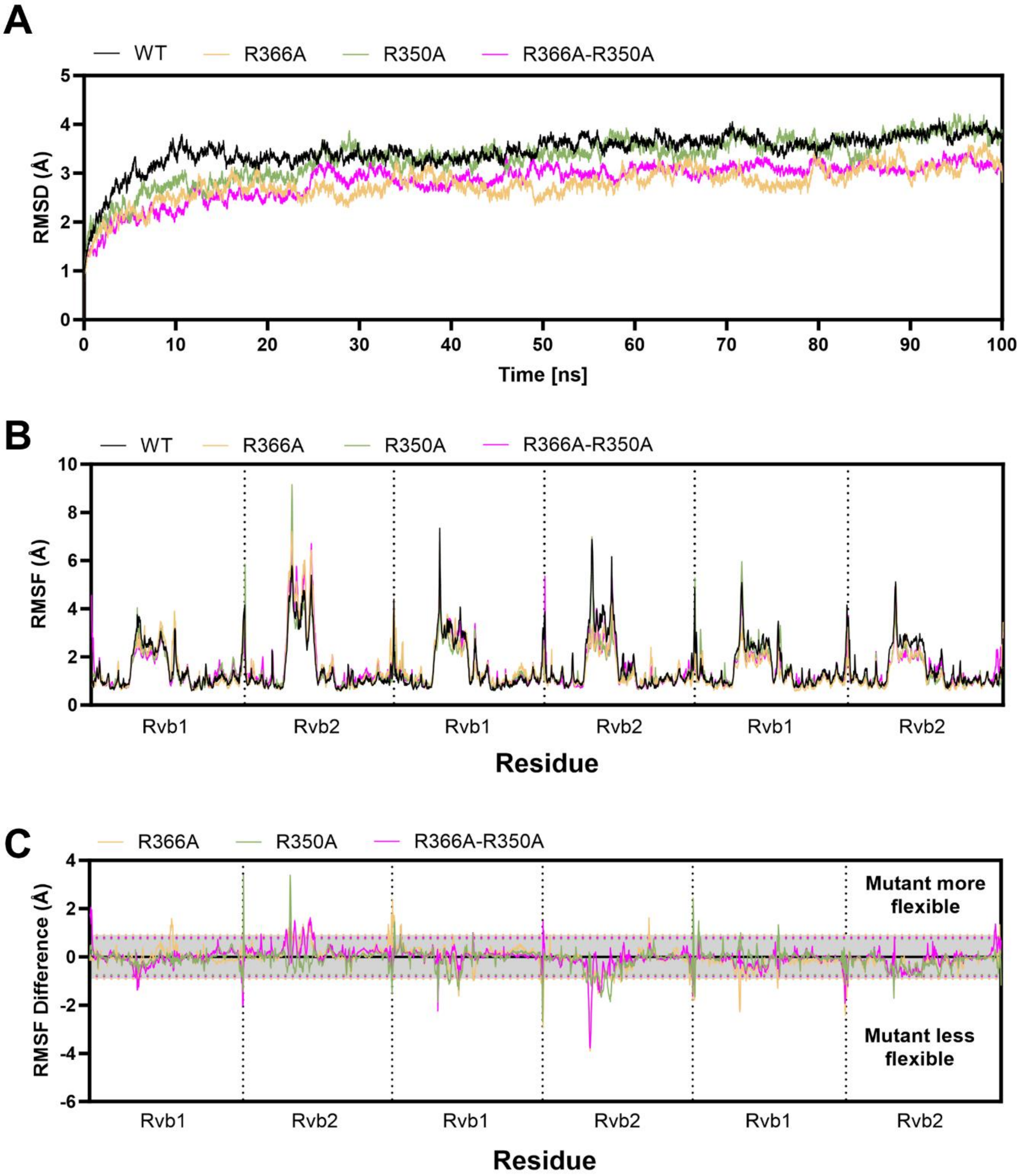
A
WT
R366A
R350A
R366A-R350A
RMSD (Å)
0
1
2
3
4
5
0
10
20
30
40
50
60
70
80
90
100
Time [ns]
B
WT
R366A
R350A
R366A-R350A
RMSF (Å)
0
2
4
6
8
10
Rvb1
Rvb2
Rvb1
Rvb2
Rvb1
Rvb2
Residue
C
R366A
R350A
R366A-R350A
RMSF Difference (Å)
4
2
0
-2
-4
-6
Mutant more flexible
Mutant less flexible
Rvb1
Rvb2
Rvb1
Rvb2
Rvb1
Rvb2
Residue

### Figure S6

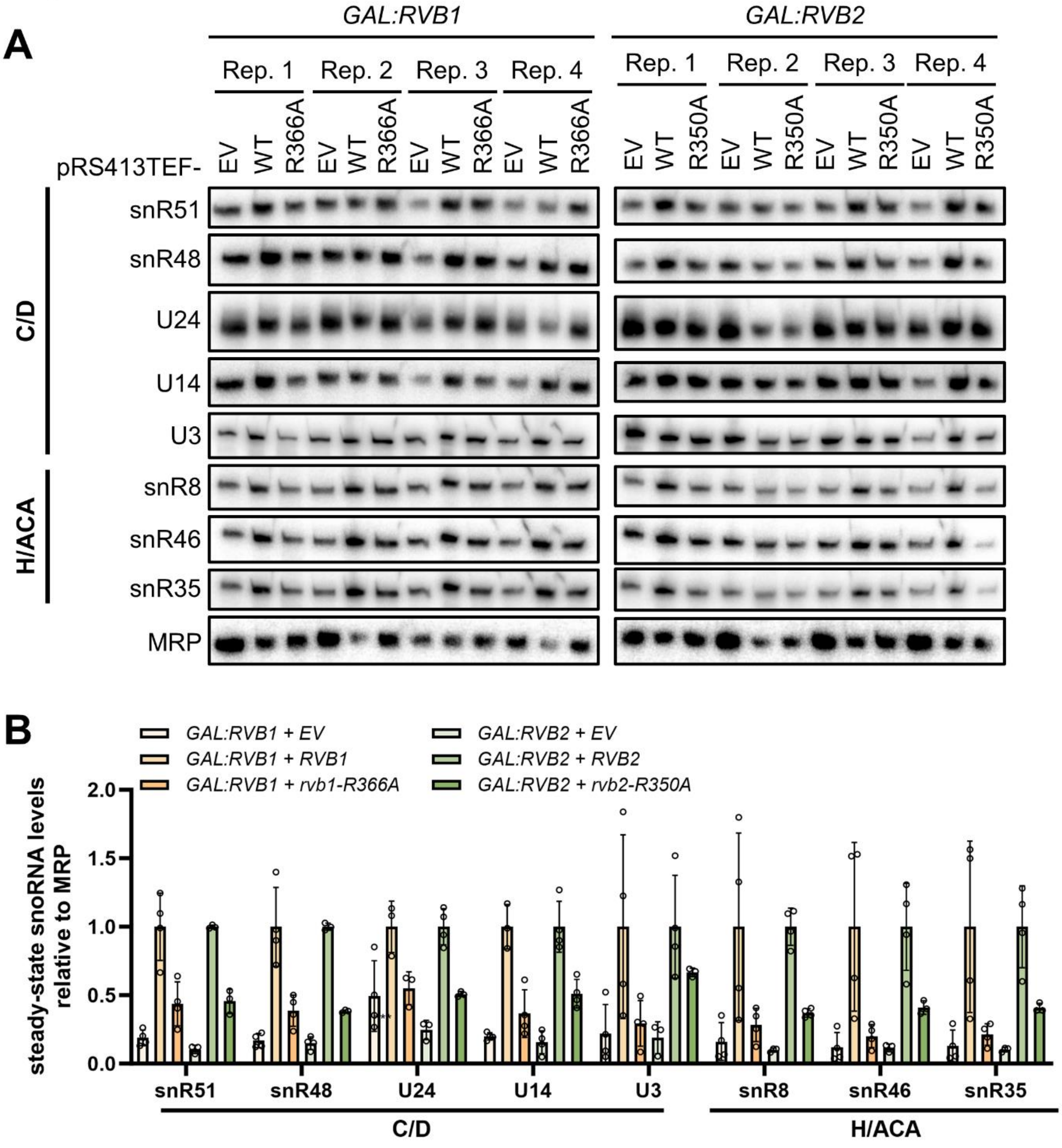

A
GAL:RVB1
GAL:RVB2
Rep. 1
Rep. 2
Rep. 3
Rep. 4
pRS413TEF-
EV
WT
R366A
R350A
C/D
H/ACA
snR51
snR48
U24
U14
U3
snR8
snR46
snR35
MRP
B
GAL:RVB1 + EV
GAL:RVB1 + RVB1
GAL:RVB1 + rvb1-R366A
GAL:RVB2 + EV
GAL:RVB2 + RVB2
GAL:RVB2 + rvb2-R350A
steady-state snoRNA levels relative to MRP
2.0
1.5
1.0
0.5
0.0
snR51
snR48
U24
U14
U3
snR8
snR46
snR35
C/D
H/ACA
